# On-Demand Seizures Facilitate Rapid Screening of Therapeutics for Epilepsy

**DOI:** 10.1101/2024.08.26.609726

**Authors:** Yuzhang Chen, Brian Litt, Flavia Vitale, Hajime Takano

## Abstract

Animal models of epilepsy are critical in drug development and therapeutic testing, but dominant methods for pharmaceutical evaluation face a tradeoff between higher throughput and etiological relevance. For example, in temporal lobe epilepsy, a type of epilepsy where seizures originate from limbic structures like the hippocampus, the main screening models are either based on acutely induced seizures in wild type, naïve animals or spontaneous seizures in chronically epileptic animals. Both types have their disadvantages – the acute convulsant or kindling induced seizures do not account for the myriad neuropathological changes in the diseased, epileptic brains, and spontaneous behavioral seizures are sparse in the chronically epileptic models, making it time-intensive to sufficiently power experiments. In this study, we took a mechanistic approach to precipitate seizures “on demand” in chronically epileptic mice. We briefly synchronized principal cells in the CA1 region of the diseased hippocampus to reliably induce stereotyped on-demand behavioral seizures. These induced seizures resembled naturally occurring spontaneous seizures in the epileptic animals and could be stopped by commonly prescribed anti-seizure medications such as levetiracetam and diazepam. Furthermore, we showed that seizures induced in chronically epileptic animals differed from those in naïve animals, highlighting the importance of evaluating therapeutics in the diseased circuit. Taken together, we envision our model to advance the speed at which both pharmacological and closed loop interventions for temporal lobe epilepsy are evaluated.

## INTRODUCTION

Epilepsy, a set of neurological disease syndromes characterized by recurrent, spontaneous seizures, is a debilitating condition that affects millions of people worldwide (*1, 2*). Even with the development of over forty anti-seizure medications (ASMs) over the course of the last century, between 15 – 30 % of patients are unable to achieve seizure freedom (*3–5*). The percentage of patients with drug resistant seizures has remained constant despite the introduction of multidrug therapies and newer ASMs classes (*6*).

The lack of screening in etiologically relevant models may be one reason for the disconnect between the increasing types of medications and the stubbornness of drug resistant seizures to treatment (*7–9*). Many patients with drug resistant seizures have temporal lobe epilepsy (TLE), a type of epilepsy where seizures originate from temporal lobe structures such as the hippocampus (*10*). The brains of TLE patients typically have structural and molecular changes, including hippocampal sclerosis, axonal sprouting, and receptor alterations (*11–14*). The resulting epileptic network functions differently from that of the healthy brain. This divergence may be why ASMs that work in a naïve, wild type animal screen are ineffective in epileptic patients.

Fortunately, TLE is well modeled in animals (*15–18*). The intrahippocampal kainate (IHK) model of epilepsy captures key markers of human TLE – animals exhibit hippocampal sclerosis, mossy fiber sprouting, and spontaneous limbic seizures (*19, 20*). Despite its lengthy history, the IHK model was only recently introduced into the Epilepsy Therapy Screening Program (ETSP) pipeline (*8, 21*). There were many factors contributing to this delay. Spontaneous seizures, by their very nature, are sparse, irregular, and unpredictable (*22–24*). Compared to testing ASMs in wild type animals by acutely inducing seizures via convulsants or kindling, testing therapeutics in etiologically relevant models is often a much more lengthy and unwieldly process.

To address the challenges of using etiologically relevant models for therapy screening, we asked whether we could harness the epileptic circuit to generate seizures on demand. Of particular interest to seizure initiation and propagation is region CA1 of the hippocampus. The CA1 is the main output of the hippocampus in the canonical tri-synaptic circuit (*25*). *In vitro*, electrical activation of excitatory inputs to the CA1 can generate seizure-like bursting (*26*). In animals, calcium imaging demonstrates that CA1 principal cells are highly active during acute convulsant induced seizures (*27*). In addition, changes in the CA1 cellular population during epileptogenesis, such as an increase in burster cells, makes the CA1 predisposed to rapid firing that can elicit epileptic events (*28, 29*). Thus, we hypothesized that hyperexcitation of CA1 principal cells would activate etiologically relevant mechanisms and initiate seizures in epileptic animals.

Specifically, we investigated whether selective optogenetic activation of CA1 principal cells could precipitate time-locked seizures in freely moving epileptic animals (Figure. 1). We used a modified version of the IHK model that preserves CA1 structure, making it more similar to human TLE. To validate the model, we first compared the induced activity to spontaneous seizures. Next, we compared seizures induced in epileptic animals to those in wild type, naïve animals to ascertain whether unique therapeutically relevant features are evident in the epileptic brain. Finally, we attempted to shut down induced seizures with known anti-seizure medications. We present evidence that hypersynchronous excitation of CA1 principal cells can induce focal to bilateral tonic-clonic seizures in mice, and that these seizures can be used to evaluate the therapeutic efficacy of both pharmacological and time-sensitive treatments.

**Figure 1:**
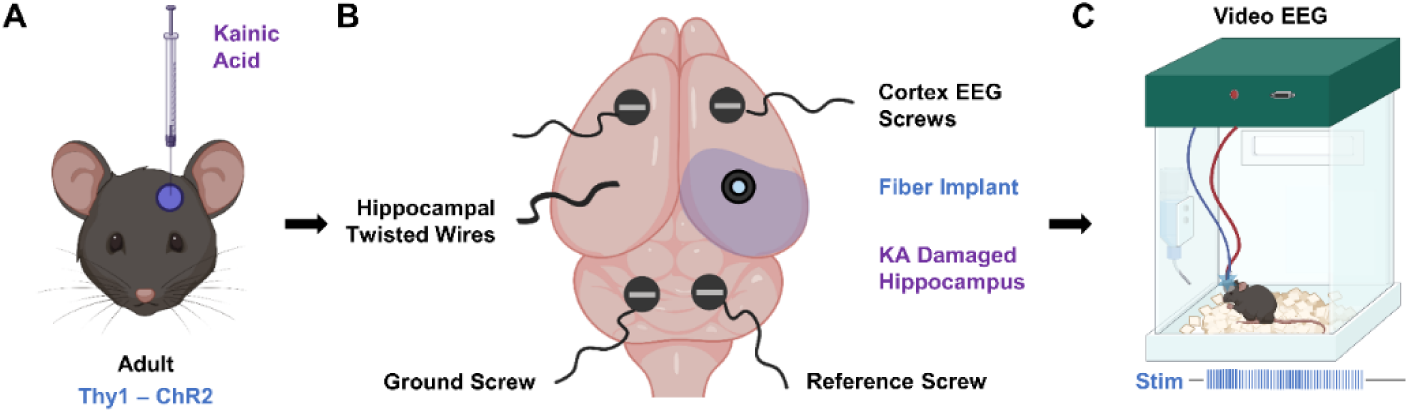
Experimental schematic for on-demand seizure induction in epileptic animals. (A) Chronic epilepsy was induced in Thy1-ChR2 mice via intrahippocampal kainic acid injection into the CA3. (B) Mice were implanted with EEG recording apparatus consisting of two cortical screws, one set of insulated braided wire targeting the hippocampus, a ground screw, and a reference screw. A fiber was positioned so that the tip illuminated the CA1. (C) 10 Hz of 473 nm light delivered into the CA1 activated Thy1-ChR2 neurons and induced seizures on demand *in vivo*.

## RESULTS

To avoid confusion and ensure clarity in the interpretation of the results presented in this manuscript, we define the following terms:

**Afterdischarge:** Afterdischarges, also referred to as induced activity or electrographic events, refer to long lasting (5s or more) electrical activity elicited following optical stimulation. This denotation includes both seizures and spikes. We use ’afterdischarges’ interchangeably with ’induced activity’.

**Seizures:** We use the term ’seizure’ specifically to describe generalized seizures that had behavioral components verified on video. Mostly these are tonic-clonic seizures with Racine Scale 3 or higher, but occasionally we observed milder seizures with Racine Scale 1 or 2.

### CA3 kainate injection preserves CA1 cell layer targeted during on-demand seizure induction

The on-demand seizure induction procedure in mice expressing excitatory channelrhodopsins in CA1 pyramidal neurons (Thy1-ChR2 mice) is a three-part process (Fig. 1). First, we induced chronic epilepsy in the mice via CA3 IHK injection (Fig. 1A). An electroencephalogram (EEG) implantation surgery followed, during which two recording screws were inserted into the cortex, and two additional screws – a ground and a reference – were inserted into the cerebellum (Fig. 1B). An implanted optical fiber targeting the CA1 ipsilateral to the IHK injection allowed for optical excitation at the seizure focus, while a braided wire targeting the CA1 contralateral to IHK injection provided information on inter-hippocampal activity (Figs. 1B & 2A middle, 2A bottom right). After the mice recovered for a week, they underwent 24-hour continuous video/EEG monitoring, during which 10 Hz, 473 nm optical stimulation was applied to induce seizures on demand (Fig. 1C).

The canonical injection site for IHK models is the dorsal CA1 (*30*); however, the destruction of the CA1 cell layer due to IHK injection is fundamentally incompatible with our approach for inducing seizures. Kainate, a glutamate agonist, induces hyperexcitation of cells in the vicinity of the injection site, leading to cell death and gliosis (*31*). The damage is clearly visible in the CA1, and the sclerosis is hypothesized to be the center of seizure generation (*32*) (Figure 2A top). Thy1 expression in the CA1 localizes to principal cells (*33*), so it is critical to preserve the health of the CA1 cell layer. To do so, we moved the IHK injection site to the medioventral CA3, which is 0.7 mm posterior, 1.4 mm lateral, and 1.6 mm ventral to the canonical CA1 injection site. CA3 IHK animals have a sparse CA1 layer (Fig. 2A middle) compared to naïve animals (Fig. 2A bottom left), while CA1 IHK animals often do not have a visible CA1 layer (Fig. 2A top).

**Figure 2:**
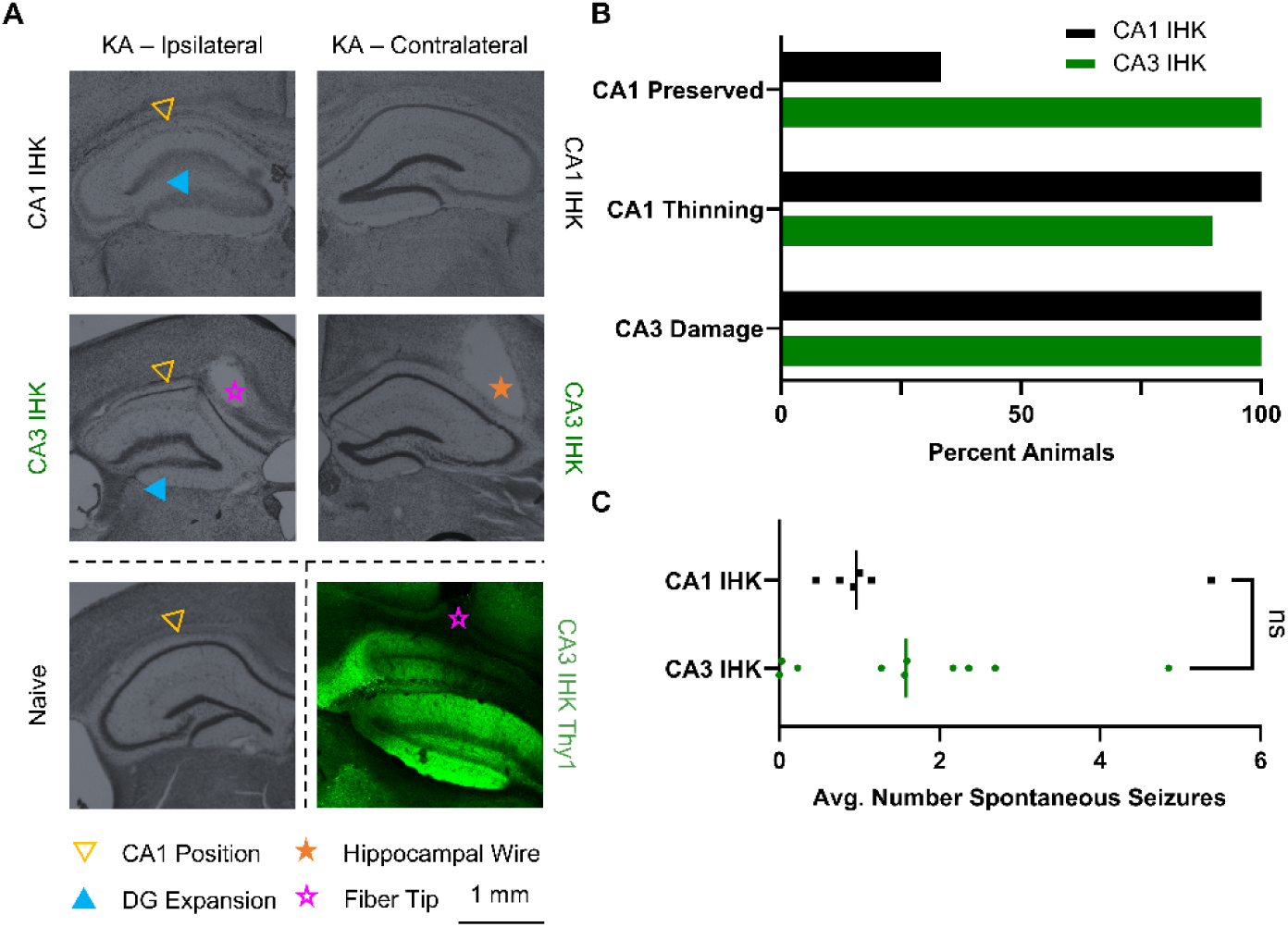
Intrahippocampal kainic acid (IHK) injection into the CA3 reduced CA1 damage without impacting average spontaneous seizure count. (**A**) Cresyl violet staining of hippocampal slices from IHK injected animals visualized IHK induced damage to the hippocampus. The CA1 layer was eliminated in CA1 IHK animals (top). In the CA3 IHK animals, the CA1 layer was present but thinned (middle). Both CA1 IHK and CA3 IHK animals displayed expansion of the dentate gyrus structure and extensive CA3 damage when compared to naïve animals (bottom left). Fiber was implanted above cells expressing Thy1-ChR2 in the CA1 (bottom right). (**B**) Two thirds (4) of CA1-IHK animals did not have a visible CA1 cell layer, while the rest (2) had a thinned cell layer. In contrast, all CA3 IHK (10) animals had a visible CA1 cell layer. All 10 CA3 IHK and 6 CA1 IHK animals had extensive damage to the CA3 coupled with expansion of the dentate gyrus (DG). (**C**) Two tailed Mann-Whitney test comparing the average number of spontaneous seizures per day showed no significant difference (p = 0.635) between CA1 IHK and CA3 IHK animals.

Comparative analysis of brain slices extracted from the CA1 IHK (n = 6) and the CA3 IHK (n = 10) mice showed greater CA1 preservation in the CA3 IHK group (Fig. 2B). Nissl staining using cresyl violet highlighted neural structures and stained the CA1 principal cell layer. Slices extracted at AP – 2.0 mm from bregma showed that the CA1 cell layer was present in only one third of CA1 IHK animals. In contrast, the CA1 cell layer was present in all CA3 IHK animals (Fig. 2B top). In both groups, the CA1 layer tended to be thinned when present. However, the CA1 in 1 CA3 IHK animal did not appear to be thinned (Fig. 2B middle). All animals in the CA3 IHK and the CA1 IHK groups exhibited CA3 damage and morphological changes in the dentate gyrus (Fig. 2A, 2B bottom). Thus, CA3 damage was not unique to the CA3 IHK group; it was already present in animals with the canonical CA1 IHK injection.

While preserving the CA1 structure was important to our model, we wanted to ensure that shifting the IHK injection site to the CA3 did not alter the number of spontaneous seizures an animal experienced per day. Thus, we acquired continuous video-EEG recording on the same CA1 IHK and CA3 IHK animals and tracked spontaneous seizures. We found that the average number of spontaneous seizures a day was similar between the two groups. The CA1 IHK animals experienced an average of 1.6 spontaneous seizures a day while the CA3 IHK animals experienced an average of 1.7 spontaneous seizures a day (Supplemental Figure S1). The nonparametric two tailed Mann-Whitney test, which compared the difference between the median daily spontaneous seizure count of the CA1 IHK (1.0) and CA3 IHK (1.6) animals, also failed to find a significant difference between the two groups (p = 0.635) (Fig. 2C).

### Optogenetically induced seizures resemble spontaneous tonic-clonic seizures in epileptic mice

In epileptic, freely moving Thy1-ChR2 mice (n = 10, 7 males, 3 females), we observed spontaneous generalized seizures, as exemplified in Figure 3A. Over one week of baseline recording, animals experienced on average 1.0 ± 1.8 (mean ± standard deviation) behavioral spontaneous seizures per day (Fig. S1A). We then optically stimulated Thy1-ChR2 expressing neurons in the CA1 ipsilateral to the IHK injection site using 10 Hz, 473 nm light several times a day at one-hour intervals. The threshold laser power and stimulation duration, defined as the minimum necessary for consistent (> 66%) induction of afterdischarges lasting a minimum of 5 s, varied between the animals, with an average threshold laser power of 9.14 ± 4.75 mW and an average stimulation duration of 6.30 ± 1.64 s (Fig. S2B, S2C). The flowchart to determine the thresholds is described in Fig. S2A. Using the threshold stimulus, activity was induced approximately 88.7 ± 8.8 % of the time in the epileptic animals (Fig. S2D). Average duration of induced activity, mainly generalized seizures with behavioral components, was 30.98 ± 4.69 s (Fig. S2E). After optical stimulation began, animals experienced 1.8 ± 3.5 spontaneous behavioral seizures per day on average (Fig. S1A). A two tailed Wilcoxon rank sum test on the distribution of daily spontaneous seizure counts before and after stimulation did not find that the difference of the two groups’ medians diverged from 0 (p = 0.1336).

**Figure 3:**
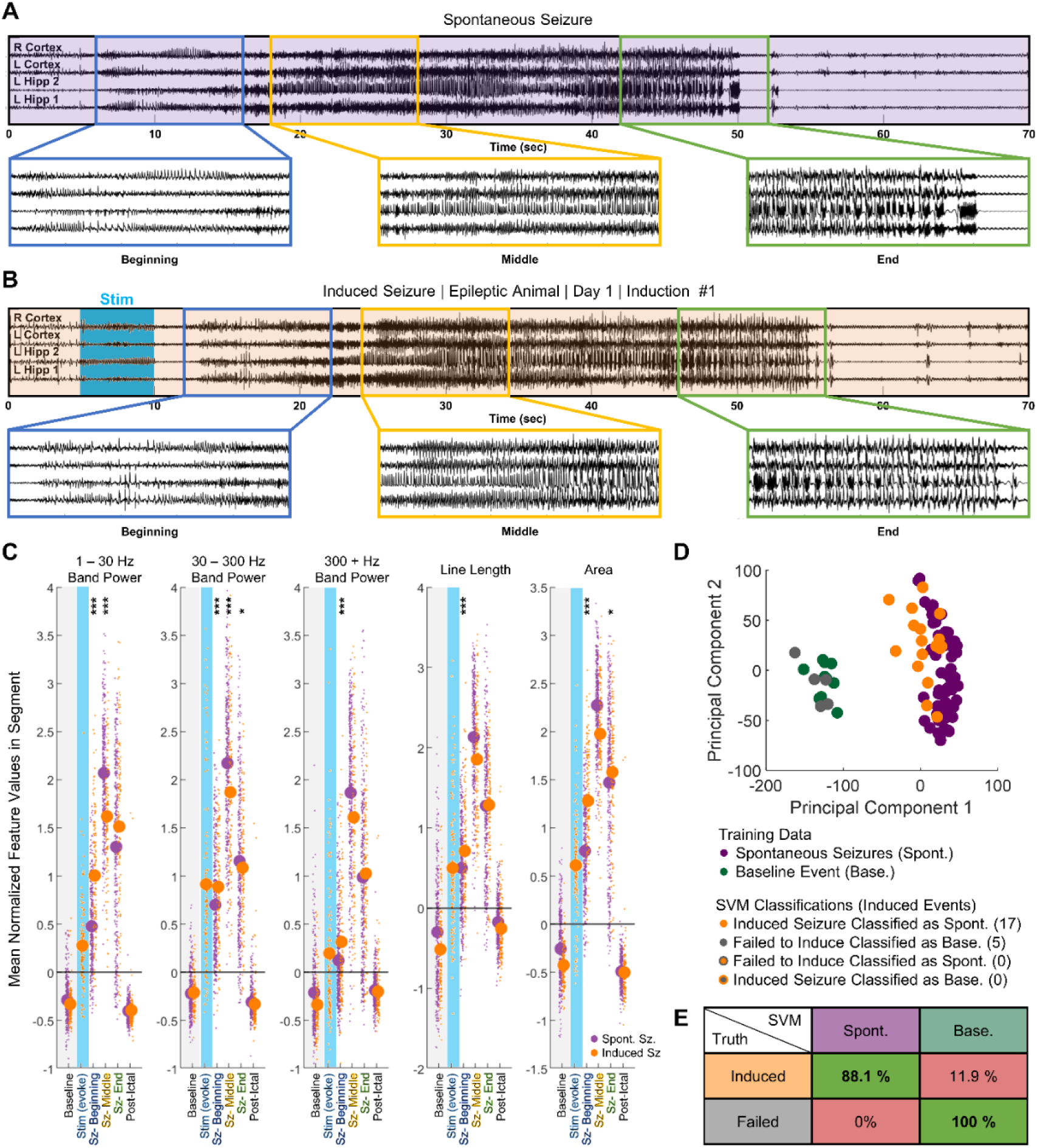
Induced seizures resembled naturally occurring spontaneous seizures in chronically epileptic animals. **(A**) Example of electrographic signal from spontaneous seizure in epileptic animal, with 3 segments of 10 s enlarged for clarity. (**B**) Electrographic signal from first induction in the same animal. (**C**) Change in features from baseline (gray) to computationally defined segments (first tercile – beginning, second tercile – middle, and final tercile – end). In the first tercile, the extent of increase in 1 – 30 Hz band power, 300 + Hz band power, line length, and area significantly differed between 138 induced (n = 10) and 337 spontaneous behavioral seizures (n = 8). Differences become less significant by the middle and the final third. Linear Mixed Effect Model: * p < 0.05, ** p < 0.01, *** p < 0.001 (**D**) Linear Support Vector Machine (SVM) classified inductions from the animal in (A) and (B). Successfully induced seizures were more closely associated with spontaneous seizures than with either baseline activity or optogenetic activations that failed to induce afterdischarges exceeding 5 seconds. (**E**) Compiled table of SVM accuracies across epileptic animals (n = 8). Induced activity was classified as similar to spontaneous seizures in 88.1% of all successful activations. Failed activations were classified as baseline 100% of the time.

We next sought to determine whether EEG signals differed between spontaneous seizures and induced seizures. First, we analyzed standard EEG features (*34*) such as band power at three different frequency ranges, line length, and area, in overlapping 500 ms sections (Fig. S3). These features were normalized to a short pre-stimulation baseline period, after which a K nearest neighbor classifier was employed to determine the seizure durations (Fig. S4). Subsequently, we divided the activity into three segments (beginning, middle, ending). This approach allows us to compare features across seizures of varying durations. Our analysis also shows that features vary widely within each animal; to account for the high intra-animal variation, we used an interactive linear mixed effect model that accounted for intra-animal residuals to identify common inter-animal trends over time. This methodology allows us to identify changes across animals, changing the independent unit in the analysis to animals instead of seizures.

Spontaneous behavioral seizures underwent stereotyped changes throughout the three segments (Fig. 3A). Visually, we noted that the beginning third of the seizure was characterized by an increase in spiking activity. The middle third was characterized by higher frequency activity and behaviorally resembled the tonic phase of a tonic-clonic seizure (*35*). The final third was characterized by bursts and behaviorally resembled the clonic phase of a tonic-clonic seizure.

Next, we quantified the feature space changes within the 337 spontaneous seizures from 8 animals using the linear mixed effect model (Fig. 3C). In the beginning third, there was a significant increase in the area (p < 1e^-4^), all band power frequencies (p < 1e^-4^), and line length (p < 1e^-4^) over baseline values. A further increase occurred in all three features (p < 1e^-4^) between the beginning and middle third of the seizure before decreasing between the middle and final third of the seizure (p < 1e^-4^). The elevated values in the final third were still significantly higher than baseline (p < 1e^-4^).

Induced activity predominantly consisted of generalized seizures (Fig. 3B). Like in the spontaneous seizures, most of induced seizures underwent three visually distinct phases. In the beginning, there was a general increase in spiking amplitude and the emergence of higher frequency activity that was occasionally accompanied by behaviors such as freezing. The middle portion resembled the tonic phase of a seizure – the animal displayed forelimb clonus, shaking, or stiffening of the tail. In the final third, bursts began to emerge on the EEG signal, and the animal entered the clonic phase of the seizure, during which it uncontrollably reared, backpedaled, jumped, or fell on its side (Supplemental Figure S5). Occasionally, optogenetic stimulus induced seizures of lower severity (Supplemental Figure S6), afterdischarges without seizure behavior, or failed to cause afterdischarges (Supplemental Figure S7). In the feature space, induced seizures followed a similar trend as spontaneous seizures (Fig. 3C). Line length, area, and band power significantly increased between baseline and the beginning third (p < 1e^-4^). A further increase occurred between the beginning and the middle third (p < 1e^-4^). Between the middle and final third, a significant decrease in area (p < 1e^-4^), line length (p < 1e^-4^), and 30 Hz + band power (p < 1e^-4^) occurred. However, unlike in spontaneous seizures, the decrease in 1 – 30 Hz band power between the middle and final third in induced seizures was not significant (p = 0.10).

We then compared the progression of spontaneous and induced seizures in the feature space. All confirmed spontaneous seizures had a Racine score of 3 or greater. To ensure a fair comparison, we used two criteria to filter the induced activity prior to comparison: 1) afterdischarges must have lasted a minimum of fifteen seconds and 2) Racine seizure score had to be greater than or equal to 3. 138 induced seizures from 10 animals fit these criteria. As shown in Fig. 3C, we used a linear mixed effect model to compare the changes in the band power, line length, and area of each segment to the baseline values. In the beginning third, the extent of increase across many features significantly differed between the induced and the spontaneous seizures. Specifically, the increase in area was significantly higher for induced seizures (p < 1e^-4^), as were the increases in line length (p < 1e^-4^), 1 – 30 Hz band power (p < 1e^-4^), 30 – 300 Hz band power (p < 1e^-4^), and 300 – 1000 Hz band power (p < 1e^-4^). By the middle third, significant differences remained in the 1 – 30 Hz band power (p < 1e^-4^) and 30 – 300 Hz band power (p = 0.0001). However, the relationship was now inverted – the 1 – 300 Hz band power increased more for spontaneous seizures than for induced seizures. The changes in other features, such as 300 – 1000 Hz band power (p = 0.3866), line length (0.7649), and area (p = 0.1962), were no longer statistically significant. In the final third, the increase in area (p = 0.0259) and 30 – 300 Hz band power (p = 0.0160) were slightly significant. The increase in 1 – 30 Hz band power was no longer significantly different (p = 0.073), and the increases in 300 – 1000 Hz band power (p = 0.3245) and line length (p = 0.1172) remained not significant.

To further quantify the similarity between spontaneous and induced activity, we trained individual one-class support vector machine (SVM) classifier to classify whether the induced activity was a seizure or not. The SVM is trained exclusively on data from spontaneous seizures and baseline activity; as such, the SVM is not designed to differentiate between induced and spontaneous seizures. Induced activity is then presented to the SVM, which classifies it as either in the category of spontaneous seizures or baseline activity. A sample classification output for one animal is shown in Fig. 3D. In this animal, all optogenetic activations that resulted in a minimum afterdischarge length of 5 s were classified as spontaneous seizures by the classifier. Meanwhile, inductions that failed to induce afterdischarges lasting more than 5 s were all classified as baseline. When we integrated the results from all 8 SVM classifiers, we found that activations that induced afterdischarges lasting longer than 5 s were classified as spontaneous seizures 88.1 % of the time, whereas 11.9 % of the time they were inaccurately classified as baseline. Meanwhile, activations that failed to induce afterdischarges longer than 5 s were classified as baseline 100 % of the time (Fig. 3E). Taken together, the results show that synchronizing CA1 principal cell activity in epileptic mice generated seizures that resembled naturally occurring spontaneous seizures, both behaviorally and on EEG, with the minor exception of EEG features during seizure onset.

### Induced activity significantly differs between epileptic and naïve animals

During epileptogenesis, neural networks in the brain undergo various changes ranging from the modification of membrane receptors to the formation of new synapses (*36–38*). We hypothesized that these changes are critical for successful on-demand seizure induction. To test our hypothesis, we attempted to optically induce seizures in naïve, wild type Thy1-ChR2 mice and compared the induced activity in naïve mice to the induced activity in epileptic mice.

Initial optical provocations in naïve animals (n = 7, 3 males, 4 females) typically caused low frequency, high amplitude spiking activity (Figure 4A). While long lasting, these afterdischarges did not have a behavioral seizure manifestation. Animals displayed independent movement, exploration, or grooming during the afterdischarges (Supplemental Figure S8A). Behavioral differences between initial naïve and epileptic inductions did not result from changes in the optical induction parameters – the naive threshold laser power (6.17 ± 1.58 mW) and stimulation duration (5.67 ± 1.03 s) both did not significantly differ from that of the epileptic animals (p > 0.05) (Fig. S2B, S2C). Rather, the observed behavior in the naïve animals was likely due to a lack of etiologically relevant changes in the healthy brain.

**Figure 4:**
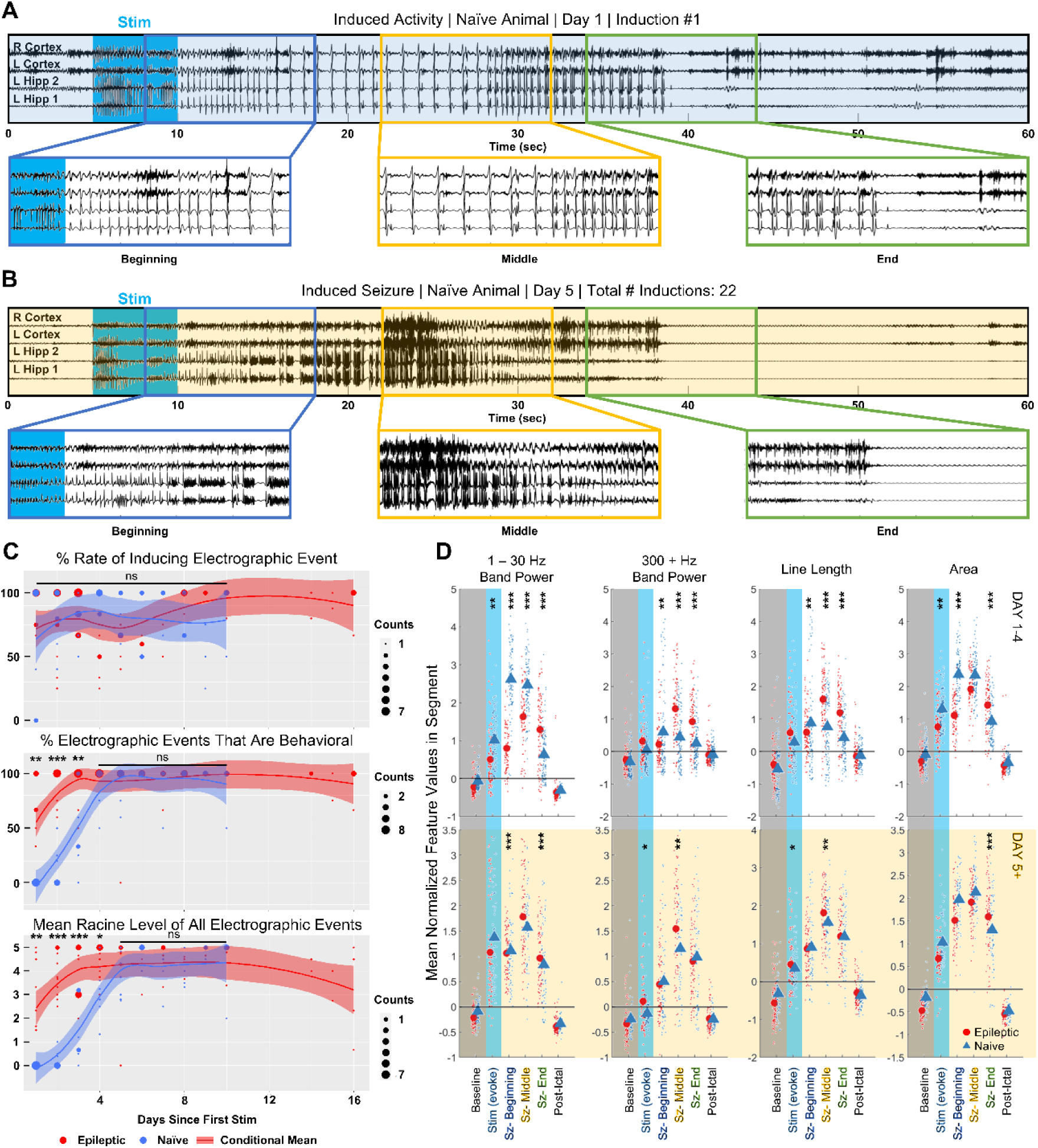
Inducing seizures in epileptic animals differs from optical kindling in naïve animals. **(A)** Initial optogenetic activations in naïve animals induced low frequency afterdischarges. Representative EEG trace with 3 segments of 10 s at the start, middle, and end of activity enlarged. **(B)** Following multiple days of stimulation, application of activation stimulus in the same animal induced Racine 5 seizures. **(C)** Rate of electrographic afterdischarges from optogenetic activation did not significantly differ between naïve and epileptic animals. Percent of behavioral electrographic events significantly differed between naïve and epileptic animals on stimulation day 1 through 3. Average Racine score of electrographic inductions significantly differed between naïve and epileptic animals on stimulation day 1 through 4. Pairwise T test: * p < 0.05, ** p < 0.01, *** p < 0.001. **(D)** In the first 4 days of stimulation, inductions in naïve (100 inductions, n = 7) and epileptic animals (87 inductions, n = 10) significantly differed in the feature space. Differences were reduced, but still existed, on stimulation day 5 or later (epileptic – 75 inductions, n = 7; naïve – 58 inductions, n = 7). Linear Mixed Effect Model: * p < 0.05, ** p < 0.01, *** p < 0.001.

After 4 consecutive days of up to 5 optogenetic activations per day at a frequency of 1 per hour, we noticed that applying the same optogenetic stimulus resulted in visibly more higher frequency activity and bursting (Fig. 4B). In addition, all 7 animals displayed stereotypical behavioral signs of seizure, including shaking, tail stiffening, rearing, wild running, and uncontrolled jumping (Supplemental Figure S8B). The increased likelihood of behavioral seizures in later inductions is reminiscent of the ’kindling model’ of epilepsy, where repeated administration of subthreshold electrical stimulation eventually causes the animal to enter a ’kindled’ state. In the kindled state, the previously subthreshold electrical stimulation can induce seizures (*39*).

To visualize the ’kindling effect’ of the optogenetic stimulation, we evaluated the rate at which both electrographic activity and seizure-like behaviors were induced over 10 experimental days (Fig. 4C top). The rate of inducing an electrographic event was defined to be the percentage of times the activation stimulus induced afterdischarges lasting a minimum of 5 s. A two-sided pairwise t test determined that, on all 10 days, the rate of inducing an electrographic event did not significantly differ between the epileptic and naïve animals (p > 0.05). After integrating data across all stimulation days, it was determined that the overall rate of inducing an electrographic event in naïve animals was approximately 86.0 ± 6.0 %, which also did not significantly differ from that of the epileptic animals (p = 0.8506, Fig. S2D). However, the average duration of induced activity in naïve animals was 20.87 ± 2.19 s, which was significantly shorter than the average duration of induced activity in epileptic animals (30.98 ± 4.69 s, p = 0.0005, Fig. S2E).

We next analyzed the percentage of electrographic events that exhibited a behavioral manifestation. This comparison between naïve and epileptic animals included behaviors such as visible forelimb clonus, uncontrolled shaking, tail stiffening, uncontrolled rearing, wild running, loss of righting reflex, and jumping (Fig. 4C middle, Table S1). We did not include behaviors in which the animal retained control over its body, such as exploring or grooming. A two-sided pairwise t test found that the percentage of electrographic events with a behavioral seizure manifestation significantly differed between the naïve and epileptic animals on the first (p = 0.0056), second (p < 1e^-4^), and third (p = 0.0088) stimulation day. No significant difference in behavioral seizure manifestations was found on stimulation day 4 or later (p > 0.05). To further quantify the behavioral differences, we compared the Racine seizure score for the induced activity (Fig. 4C bottom). A two-sided pairwise t test found that the average Racine seizure score significantly differed between the naïve and the epileptic animals on the first (p = 0.0087), second (p < 1e^-4^), third (p < 1e^-4^), and fourth (p = 0.024) stimulation day. No significant difference in Racine seizure score was found on stimulation day 5 or later (p > 0.05).

Later activations and their differing behavior indicated that the animal entered a distinct state after multiple days of optical activation. Thus, we decided to perform separate feature space comparisons of the electrographic activity between naïve animals and epileptic animals for the initial activations (day 1 – 4) and the later activations (day 5 +) (Fig. 4D). To standardize the comparison between naïve and epileptic induced activity, we followed a similar procedure to the analysis did in Figure 3. We used an interactive linear mixed effect model that accounts for the intra-animal variability to uncover the inter-animal changes in electrographic feature over three segments. The terciles of the 308 inductions in naïve animals were computationally determined by the same k-nearest neighbor classifier used in epileptic animals. To reduce noise from shorter events, only induced activity lasting more than 15 s were compared. The initial activations group was composed of 100 events from 7 naïve animals and 87 events from 10 epileptic animals. The later activations group was composed of 75 events from 7 epileptic animals and 58 events from 7 naïve animals.

Consistent with visual observation of initial activations, increases in the electrographic features versus baseline significantly differed between naïve and epileptic animals (Fig. 4D top). Overall, naïve induced activity tended to have higher area and more low frequency band power while epileptic induced activity tended to have more high frequency band power and higher signal complexity. The linear mixed effect model found significant differences in the increase of 1 – 30 Hz band power from baseline to the beginning third (p < 1e^-4^), middle third (p < 1e^-4^), and final third (p < 1e^-4^). The difference in the 300 – 1000 Hz band power from baseline was also significant between the two groups at the beginning third (p = 0.002), middle third (p < 1e^-4^), and final third (p < 1e^-4^). A similar case existed for line length at the beginning third (p = 0.008), middle third (p < 1e^-4^), and final third (p < 1e^-4^) and for area at the beginning third (p < 1e^-4^) and final third (p < 1e^-4^). The increase in area from baseline to the middle third was barely insignificant (p = 0.059).

Differences in the feature space were reduced in later activations (Fig. 4D bottom). The differences in the increase of the 1 – 30 Hz band power between the two groups were reduced in the beginning third (p = 0.0002), middle third (p = 0.051) and final third (p < 1e^-4^). The same applied to 300 – 1000 Hz band power (beginning third p = 0.60, middle third p = 0.029, final third p = 0.097), line length (beginning third p = 0.21, middle third p = 0.002, final third p = 0.11), and area (beginning third p = 0.15, middle third p = 0.56, final third p < 1e^-4^). Despite reductions between feature values in the later activations group, not all significant differences between the naïve and epileptic animals were eliminated. The persistent differences highlighted by the linear mixed effect model show the importance of evaluating novel ASMs in epileptic animals.

### Common anti-seizure medications could suppress induction of electrographic and behavioral seizures

To determine the utility of the induced seizure model for evaluating ASMs, we tested whether FDA approved and commonly prescribed ASMs, such as diazepam and levetiracetam (*40*), could reduce the rates of both electrographic afterdischarge induction (electrographic activity lasting more than 5 seconds, including seizures) and behavioral seizure (generalized seizure with behavioral component) induction in epileptic animals. Diazepam, a benzodiazepine that interacts with GABA receptors, is commonly prescribed for first line control of convulsive seizures (*41*), whereas levetiracetam is an anticonvulsant that reduces overall network excitability through effects mediated by the synaptic protein SV2A (*42*).

The experimental paradigm is shown in Figure 5A. Prior to inducing activity with the optogenetic stimulus, the baseline spontaneous seizure frequency was recorded in 9 epileptic animals. The next 2 – 5 days were used for determining and validating the consistency of the threshold stimulus. On each day of ASM testing, optogenetic activation was performed hourly to establish the drug-free electrographic discharge induction and behavioral seizure induction success rate. Then, the animal received either a single intraperitoneal injection of 800 mg/kg levetiracetam or a single subcutaneous injection of 5 mg/kg of diazepam. The activation stimulus was applied to the animal approximately 10 minutes after ASM administration and was re-applied hourly. A washout period of 48 hours, if applicable, occurred between subsequent ASM administrations. After completion of drug testing, animals underwent a post-stimulation recording period of up to 7 days.

**Figure 5:**
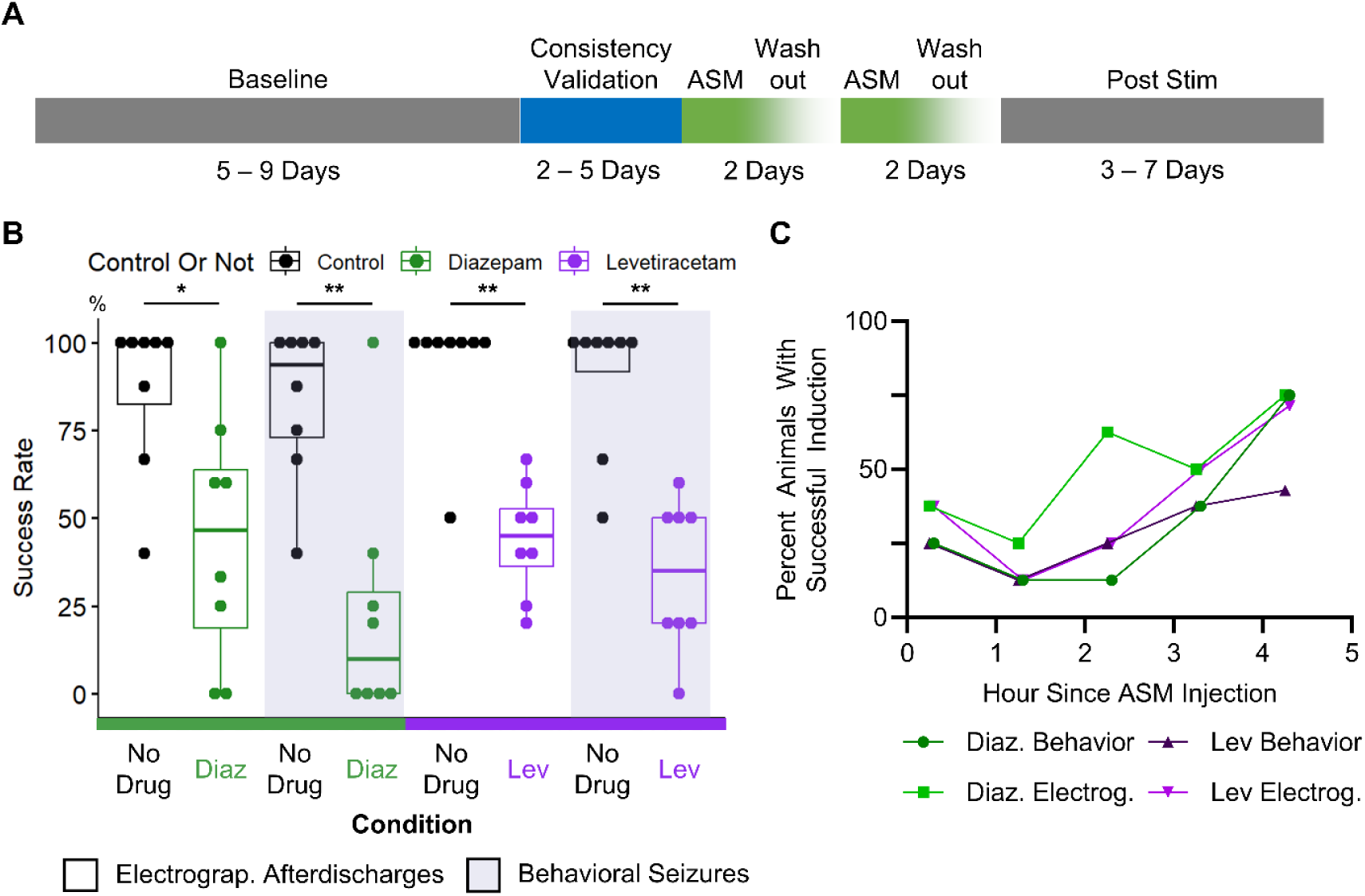
Induced seizures in epileptic animals responded to both diazepam and levetiracetam. **(A)** Experimental timeline for testing ASM efficacy in on-demand seizure model. Stimulus for seizure induction was tested for consistency before ASM application. A 48-hour washout occurred between subsequent ASM applications. **(B)** Post ASM application, rates of inducing electrographic afterdischarges (any electrographic activity lasting more than five seconds) and behavioral seizures (generalized seizures with seizure behaviors) were significantly reduced. Paired One Tailed Wilcoxon Signed Rank Test, * p < 0.05, ** p < 0.01 **(C)** Probability of successful induction of activity (averaged across all epileptic animals) increased the more time has passed since ASM injection into the mouse.

Application of diazepam significantly reduced the rates of electrographic discharge and behavioral seizure induction in 8 assessments conducted on 6 epileptic animals (Fig. 5B). Typically, after diazepam administration, optogenetic activation did not induce electrographic activity. Even when afterdischarges were successfully induced, animals typically did not display seizure-like behaviors. In all animals, the severity of behavioral seizures is reduced after diazepam administration (Supplemental Figure S9A). Integrating data from all animals, prior to drug administration, electrographic discharge induction rate averaged 86.77 ± 22.3 % and the behavioral seizure induction rate averaged 83.65 ± 21.9 %. The paired one-tailed Wilcoxon signed rank test found that after drug administration, the electrographic discharge induction rate was significantly reduced to 44.17 ± 35.8 % (p = 0.016) and the behavioral seizure induction rate was significantly reduced to 23.12 ± 34.5% (p = 0.008). One epileptic animal was a diazepam non-responder on one of two testing days. Excluding the diazepam non-responder, no tonic-clonic seizures (Racine > 3) were induced 3 hours after drug administration in any of the 5 epileptic animals. Even in the diazepam non-responder, the Racine scale of induced seizures was lower after diazepam administration (Fig. S9A).

Application of levetiracetam significantly reduced the rates of electrographic discharge and behavioral seizure induction in 8 assessments on 5 epileptic animals (Fig. 5B, S9B). After levetiracetam administration, electrographic inductions were typically unsuccessful (Supplemental Figure S10A). In mice, levetiracetam has a short half-life of approximately 3.2 hours (*43*). In our study approximately 4 hours after the injection, tonic-clonic behavioral seizures (Racine > 3) were successfully induced in 2 out of 5 epileptic animals. Integrating data from all animals, prior to drug administration, average electrographic discharge induction rate was 93.75 ± 17.7 % and the average behavioral seizure induction rate was 89.58 ± 19.8%. A paired one-tailed Wilcoxon signed sum test found that after drug administration, electrographic discharge induction rate was significantly reduced to 43.96 ± 16.1 % (p = 0.004), and the behavioral seizure induction rate was significantly reduced to 33.75 ± 21.3 % (p = 0.004). Additionally, in all animals, seizure burden, measured as the severity of behavioral seizures decreased after levetiracetam administration (Fig. S9B).

Quantifying the seizure induction rates after drug administration by hour showed that both drugs had maximal effect between the first and the second hour (Fig. 5C). As the drug was metabolized or excreted, the rate of successful seizure induction increased, eventually approximating 75% of all animals by the fourth hour. This was expected, as seizure induction rates should gradually return to pre-drug levels as the drug was cleared from the animal body. Taken together, these results showed that the induced seizure model has potential for evaluating ASM efficacy in preventing seizures and guiding calculations on ASM pharmacokinetics.

## DISCUSSION

In this study, we showed that selective activation of CA1 principal cells could induce on-demand seizures in both chronically epileptic and naïve animals. Induced seizures were reliably generated, reproducible, and bore striking similarities to spontaneously occurring seizures.

Induced activity differed between epileptic animals and naïve animals, indicating that the epileptic network is critically different from that of the healthy brain. Finally, induced seizures could be suppressed with standard ASMs, proving the model has utility for assessing the effectiveness of potential therapies.

### Seizure induction takes advantage of mechanistic changes in the hippocampus

The seizure induction procedure took advantage of naturally occurring changes in the epileptic brain to elicit seizures on demand, suggesting that hyperexcitation of principal cells could be a mechanism by which activity propagates out from the hippocampus and generalizes into behavioral seizures. The profound difference in the induced activity between naïve and epileptic animals further suggested that the epileptic circuit is a key facilitator of seizure generalization – only in the epileptic circuit could we provoke behavioral seizures from stimulation day 1.

Previous research suggests that changes in the CA1 during epileptogenesis contributes to increased bursting activity in chronically epileptic animals but stops short of establishing a causal relationship between hyperactivity and generalized seizures (*26, 28, 44, 45*). In the pilocarpine model of chronic epilepsy, CA1 principal cells in epileptic animals were prone to firing synchronous bursts of spikes when excited with inputs that induced only single spikes in naïve animals (*46*). Inputs to the CA1 also change during epileptogenesis. The temporoammonic pathway, a direct connection from the entorhinal cortex to the CA1, switched from being a highly regulated, weak excitatory input in naïve animals to a powerful excitatory pathway in epileptic animals (*47, 48*). Electrical activation of the temporoammonic pathway was also sufficient to generate bursting activity in chronically epileptic brain slices (*48*). Despite these pioneering studies, there was no direct investigation into whether seizures in the epileptic brain can initiate purely from CA1 hyperexcitation. In our work, we answer this question by showing that we can harness CA1 principal cells to induce generalized seizures on demand. We also show that these seizures are similar to spontaneous seizures in the EEG feature space; further suggesting that, in epileptic animals, this pathway is potentially active in seizure generation.

Behaviorally, induced seizures resembled tonic-clonic seizures in human patients (*49*). Induced seizures were characterized by an initial tonic phase, where the dominant behavior was freezing, and the EEG showed continuous fast spiking activity. The tonic phase is followed by a clonic phase, where the animal displayed uncontrolled movements, such as backpedaling, rearing, and jumping. During the clonic phase, we observed clearly defined bursting and large amplitude single spikes. The gross similarity in the EEG and the behavior posits a question – could CA1 hyperexcitation be one of the mechanisms by which tonic-clonic seizures naturally start? Further studies with this model could answer questions about the initiation and the termination of such seizures, with clinical implications in both the epidemiology and the treatment of medication resistant seizures.

### Application of model to testing multiple classes of ASMs, closed loop neuromodulation therapies, and studying cellular interactions

In this work, we found that seizures induced in epileptic animals could be blocked by applying two commonly used ASMs – diazepam and levetiracetam. These two medications were chosen because they act through different mechanisms – diazepam enhances GABA receptor efficacy and levetiracetam disrupts synaptic transmission (*41, 42*). Diazepam historically performs well in many seizure models. However, levetiracetam is unique among many ASMs in that it was originally missed by the standard ASM screens of the maximal electroshock seizure and subcutaneous pentylenetetrazol seizure tests. Its antiepileptic effect was only discovered using a kindling model of epilepsy, which has since been integrated into the ETSP pipeline (*50, 51*) In our model, levetiracetam had a clear anti-seizure effect. We believe that the etiological relevance of our model allows levetiracetam to exert its effect on synaptic transmission and prevent seizure generalization. These experiments show that our induced seizure model could be used for testing multiple classes of ASMs.

Outside of pharmacologics, our model could be applied to evaluating time-dependent, closed loop treatment paradigms. The model allows for precise timing of seizure onset; thus, experimenters could rapidly test the effects of both open and closed loop electrical stimulation on seizure initiation, propagation, and termination. We envision this model to enable quick advances in determining optimal targets and tuning parameters for closed loop stimulation. There is an urgent need to explore the parameter space for closed loop seizure control, as the multitude of options makes it difficult to quickly optimize responsive neurostimulators for maximal seizure control (*52*). A reliable, physiologically realistic on-demand seizure model could dramatically accelerate this process.

Alternatively, our approach can be harnessed to answer questions of biological and clinical significance. For instance, adaptation of our approach into rare disease models of epilepsy, such as the Scn1a mouse model used to study Dravet Syndrome (*53*), could produce fresh insights into how seizures spread in different models. Comparing seizure propagation across various models will also help elucidate how genetic mutations and cellular abnormalities influence seizure mechanisms across the heterogeneity of human epilepsy. Furthermore, circuit specific investigations can be conducted by integrating our model with cell-specific optogenetic or chemogenetic approaches (*54*). By evaluating seizure induction rate while suppressing or activating certain neural populations, one can quickly assess the potential of the circuit specific intervention strategy to control seizures.

Our experiments suggest that interfering with activity from the CA1 region may be one way to stop seizure progression. Diazepam and levetiracetam both reduced the likelihood of induced activity generalizing beyond the hippocampus, resulting in lower rates of electrographic activity induction and lower rates of behavioral seizure induction. As the CA1 is the main output of the hippocampus, deciphering ways to prevent hippocampal activity from generalizing could be impactful for clinical care. If abnormal hippocampal outputs could not leave the hippocampus, patients with seizures that originate in the hippocampus might not experience the debilitating effects of grand mal seizures.

### Comparison to other seizure models used in pharmacologic screening

Other models that acutely induce seizures exist, but they either do not use chronically epileptic animals or do not have as rapid onset as the one presented in this work (*30*). Some models utilize a kindling paradigm, requiring multiple days of successive activation before seizures can be reliably induced (*39*). Other approaches are performed under anesthesia (*55*) or utilize naïve animals (*56*), both of which could introduce differences in neural activity when compared to studies in freely moving epileptic animals. Overall, prior studies we reviewed do not sufficiently address the challenges of efficiently evaluating epilepsy treatments in etiologically relevant models.

Conversely, our model allows for more robust powering of studies in etiologically relevant models. We target a potential seizure mechanism in the CA1 to reliably elicit seizures hourly in freely moving animals. Our model also generates both long afterdischarges and tonic-clonic behavioral seizures. This is unlike models that rely on continuous stimulation throughout the seizure to generate behavioral events or models that pre-screen for animals that are predisposed to have spontaneous seizures (*56, 57*). In our model, all animals are used, and ictal activity naturally propagates out of the hippocampus and evolves into generalized tonic-clonic seizures. Our model is also unlike traditional kindling models because we target specific cellular populations instead of using non-specific electrical stimulus (*39*). Thus, seizures from our model are likely to be highly relevant for testing clinical treatments and developing novel approaches for epilepsy.

Long term, we envision this model could be integrated into a drug screening pipeline such as the Epilepsy Therapy Screening Program (ETSP) (*8*). Currently, the ETSP utilizes a mixture of models, including acutely induced seizures in naïve animals and spontaneous recurrent seizures in chronically epileptic, IHK rodents. An intermediate step in the ETSP evaluates how ASMs affect hippocampal paroxysmal discharges (HPDs), which are frequent non-behavioral, focal electrographic seizures, to quickly produce dose-response curves with high confidence (*58*).

However, HPDs are not generalizable across species – they are specific to the mouse model (*59*). In addition, it is unclear whether medications that prevent HPDs would also be effective against generalized convulsive seizures or partial non-convulsive seizures in humans.

In line with modern medicine’s aim to develop etiology-specific drugs as part of a precision medicine approach, it is crucial to incorporate evaluation of convulsive seizures into the screening process to discover drugs that can effectively stop these seizures. Our model is well positioned to supplant HPDs in the existing pipeline. First, on demand seizures do not require more time to prepare than HPDs. In the preparation phase, both methods require chronic epilepsy induction and an EEG implantation surgery. In the screening phase, on demand seizures produce significant time savings, as seizures can be evaluated immediately instead of waiting for HPD characterization and analysis. Furthermore, as on demand seizures are behavioral seizures, we predict that our model will respond to more categories of ASMs than HPDs, serving as a quick initial check as to whether the drugs’ actions are sustained on seizures in the epileptic brain.

Some steps are still needed prior to integration - additional scaling of the model and validation with more ASMs are critical further studies. These experiments could better characterize the model and help determine whether the model can serve as a testbed for pharmacologics that specifically target refractory, drug resistant seizures.

In all, we present a biologically relevant, higher throughput on-demand seizure model that can be used to evaluate novel pharmacologics and time-sensitive treatment paradigms. This model takes advantage of a biologically relevant mechanism to generate long afterdischarges and behavioral tonic-clonic seizures. With this model, we envision immediately increasing the speed of discovering and evaluating new clinical treatments for epilepsy and the identification of new mechanisms behind the initiation of tonic-clonic seizures.

## MATERIALS AND METHODS

### Study Design

The purpose of this study was to create a higher throughput model for evaluating ASMs in the diseased, epileptic brain. To do so, we tested whether specific optical activation of CA1 principal cells could induce generalized behavioral seizures in epileptic animals. Next, we compared these induced seizures to spontaneous seizures to establish the model’s face validity. To show that successful induction was dependent on changes in the epileptic brain, we attempted to provoke seizures in naïve animals. Comparing the induced activity between naïve and epileptic animals allowed us to establish the model’s construct validity. Finally, we attempted to use known ASMs to stop seizures. Diazepam and levetiracetam reduced likelihood of seizure induction, establishing that our model has predictive validity. Blinding was not utilized in this study.

### Animal Experiments

All procedures are approved in accordance with the lab’s Institutional Animal Care and Use Committee (IACUC) protocol at the Children’s Hospital of Philadelphia. More details in supplemental materials and methods.

### Epilepsy Induction

C57BL/6J-Thy1-ChR2-YFP mice (Jax #007612) underwent intrahippocampal injection of 50 nL, 20 mM kainic acid (Hello Bio) under anesthesia (coordinates: right CA3 AP – 2.7, ML + 3.0, DV – 3.2, right CA1 AP – 2.0, ML + 1.6, DV – 1.6). After onset of status epilepticus, 5 mg/kg diazepam was administered to reduce severity of seizures. Animals that died from epilepsy induction were excluded.

### EEG and Fiber Implantation

Animals were implanted with A-M Systems Model 1700 adapters connected by wire to two cortical screws (AP + 0.5, ML ± 1.7), two cerebellar screws (AP – 5.5, ML ± 2.0, ground and reference), and a pair of twisted hippocampal depth wires (AP – 2.0, ML – 1.7, DV – 1.6). 400-micron diameter optic fiber cannula (RWD Life Science) was positioned at 2.0 mm (AP – 2.0, ML + 1.7) depth illuminating the damaged CA1. Dental cement (Lang Dental) secured the entire apparatus.

### Video EEG and Optogenetic Stimulus

Animals were placed into a Plexiglass cage and connected to a Stellate Harmonie (Natus) recording system. A Master 8 controller that was set to 10 Hz, 25 ms pulses was used to control delivery of light from a Laserglow 473 nm laser. Laser power and stimulation duration depicted in Figs. S2B and S2C. Stimulation was performed every 1 – 3 hours over many days. Animals that had no light response at all – not even a stimulation artifact – were excluded.

### Pharmaceuticals

On each testing day, a few optogenetic activations occurred prior to drug injection to determine the pre-drug induction success rate. Animals then received either one subcutaneous injection of 5 mg/kg diazepam (Dash Pharmaceuticals) or an intraperitoneal injection of 800 mg/kg levetiracetam (Sigma Aldrich). Up to six more activations occurred every 1 – 1.5 hours following ASM administration. One tailed Wilcoxon signed rank test was used to compare success rate before and after ASM administration.

### Perfusion and Histology

Animals were deeply anesthesized and transcardially perfused with PBS and fixed in 4% paraformaldehyde in PBS. Cryoprotected brains were sliced and then stained with 0.1% cresyl violet (Sigma Aldrich) using the Nissl staining protocol. Slices were imaged on a Leica DMIRB microscope using the LasX software.

### Statistical Analysis & Computational Processing

All code is available on Github.com at the following repository: https://github.com/yuzhangc/Evoked_Seizures

#### **A.** Preprocessing

EEG was filtered with a second order notch filter to remove 60 Hz line noise. The signal was then z-score normalized. A 4 Hz 6^th^ order Butterworth high pass filter subsequently removed low frequency artifacts.

#### **B.** Feature Calculation

Features were extracted from 500 ms windows with 250 ms displacement. Full formula in supplements. Features calculated include line length, area, energy, zero crossing around mean, root mean squared amplitude, skewness, approximate entropy, Lyapunov exponent, phase locked high gamma, magnitude squared coherence, mean absolute deviation, and band power at three frequency ranges (1 – 30 Hz, 30 – 300 Hz, 300 + Hz). Features were z-score normalized for machine learning.

#### **C.** Induced Activity Length Determination and Calculation of Thirds

An unsupervised K nearest neighbor model was trained with z-score normalized input from one spontaneous seizure. The model had three output classes – two were classified as ‘seizure’ and the last one was ‘baseline.’ The model was then used to determine induced activity length across all events from all animals. Two tailed Wilcoxon rank sum test and/or two-sided pairwise t test was used to compare induction success rates. Wilcoxon rank sum test was used to compare the threshold duration and the threshold power distribution of the naïve and epileptic animals.

Features calculated in Section B were then segregated into 1 of 6 time points based on the length of induced activity: 1) before stimulation, 2) during stimulation, 3) initial/beginning third, 4) second/middle third, 5) final/ending third, and 6) post stimulation (30 seconds). Linear mixed effect models (using the lmer function in lmerTest package) was used to determine how the comparison (such as epileptic vs naive or spontaneous vs induced) influenced the EEG features over time points.

#### **D.** Spontaneous Seizure Detection and Video EEG Behavioral Scoring

Spontaneous seizures were detected using custom-written Matlab code (*60*). Comparisons on daily spontaneous seizure frequency used a two tailed Mann Whitney test. Uncontrolled behaviors and the Racine scale (*61*) associated with the behaviors are as follows, and described in Table S1: (1) – clear freezing / flattening of the body, (2) – tail stiffening, forelimb clonus, uncontrolled shaking, (3) – rearing, (4) – wild run, and (5) – uncontrolled jumping, loss of righting reflex. If no behavior or movement was displayed, the score was 0. Behavioral manifestation was compared with a two sided pairwise t test.

#### **E.** Support Vector Machines

A new support vector machine for classifying induced activity was trained per animal. EEG features in (B) were calculated for baseline segments for training unique SVM classifiers. The baseline features were combined with spontaneous seizure features to form training inputs.

Testing data consisted of features from induced activity. Accuracy and error rates were calculated based on ground truth, where a successful induction was defined as an electrographic event with afterdischarges lasting a minimum of 5 seconds.

## Acknowledgments

We acknowledge Dr. Douglas Coulter for his invaluable scientific insights and support on our project. We acknowledge the support of Alicia White and Emily Schellinger for their technical assistance in this project, as well as Dr. Srdjan Joksimovic and Dr. Anthoni Goodman for their helpful suggestions. We also acknowledge Dr. Mary Putt, Director of the IDDRC Biostatistics and Data Science Core, for her statistical advice and consultation.

## Author contributions

Conceptualization: YC, FV, HT, BL Methodology: YC, FV, HT Investigation: YC, HT Visualization: YC, HT

Funding acquisition: BL, FV, HT Project administration: FV, HT, BL Supervision: HT, FV, BL

Writing – original draft: YC

Writing – review & editing: YC, HT, FV, BL

## Data and materials availability

Code is available on Github at the following repository: https://github.com/yuzhangc/Evoked_Seizures. Raw data and video clips are available upon request by contacting the corresponding author.

