## Supplemental for "On-Demand Seizures Facilitate Rapid Screening of Therapeutics for Epilepsy"

### Supplementary Materials:

List of Supplementary Materials

- Materials and Methods
- Table S1
- Figs. S1 – S 10

### Materials and Methods

#### Animals

All procedures are approved in accordance with the lab's Institutional Animal Care and Use Committee (IACUC) protocol at the Children's Hospital of Philadelphia which is an Association for Assessment and Accreditation of Laboratory Animal Care International (AAALAC) accredited site.

Adult (12 – 20 weeks old) heterozygous C57BL/6J-Thy1-CHR2-YFP mice were bred by crossing homozygous C57BL/6J-Thy1-ChR2-YFP mice (Jax #007612) with either wild type C57BL/6J mice, homozygous B6/J-PV-IRES-Cre (Jax # 017320) or heterozygous B6N/J-SST-IRES-Cre (Jax #018973) strain.

#### Chronic Epilepsy Induction

Chronic epilepsy was induced by intrahippocampal kainate injection into region CA3 or region CA1. To induce anesthesia, mice received subcutaneous injection of buprenorphine (0.5 mg/mL) and 1 – 2 % isoflurane in oxygen. A hole through the skull was drilled above the right CA3 (AP – 2.7, ML + 3.0) or the right CA1 (AP – 2.0, ML + 1.6). 50 nL of 20 mM kainic acid (Hello Bio) was injected into the medioventral CA3 (DV – 3.2) or the dorsal CA1 (DV – 1.6) over the course of 50 seconds using a Drummond Nanoject III. Seizures usually began within 20 minutes of injection and animals experience four or more seizures in the first hour after injection. During seizures, animals typically exhibited circular movements, freezing, wild run, digging, and shaking. Within one hour after the onset of status epilepticus, diazepam (5 mg/kg) was administered subcutaneously to terminate and/or reduce the severity of seizures. Animals that died from KA treatment were excluded from the study. Studies were conducted after a minimum of three weeks post chronic epilepsy induction.

#### EEG and Optical Fiber Implantation Surgery

Animals underwent a simultaneous EEG adapter and optical fiber implantation surgery. First, 4 exposed silver wires (A-M Systems) and a twisted insulated wire (PlasticOne) were soldered to the A-M Systems Model 1700 adapter and secured with epoxy. Then, holes were drilled into the mouse skull at the following positions: Bregma AP – 5.5, ML – 2.0 (Reference); Bregma AP – 5.5, ML + 2.0 (Ground); Bregma AP + 0.5, ML + 1.6 (Right Cortex); and Bregma AP + 0.5, ML – 1.6 (Left Cortex), and four stainless-steel screws (JJ Morris) were inserted. Another hole was drilled at Bregma AP – 2.0, ML + 1.7 (Right CA1) for positioning 2 mm deep, 400 micrometer diameter optic fiber cannula (RWD Life Sciences). At Bregma AP – 2.0, ML – 1.7, a hole was drilled and twisted wires were lowered until they were at a depth of – 1.6 mm (contralateral CA1). Then, the four wires on the adapter were connected to the EEG screws. Dental cement was used alongside superglue to secure the entire apparatus to the skull (Lang Dental).

#### Video EEG Setup and Optogenetic Stimulus for Seizure Induction

For video EEG recording, animals were placed into a custom-made Plexiglass cage. FC-LC fibers (RWD) were attached to the optic cannula and a flexible cable (A-M Systems) connected the headcap assembly to a commutator (PlasticsOne SL12C), allowing the mouse free movement during recordings. Concurrent video-EEG (2 kHz sampling rate) was acquired using a Stellate Harmonie system (Natus Medical). Baseline recording was performed for at least 72 hours before optical stimulation. During the entire recording, water and food were provided ad libitum. Mice were in a 12-hour light/dark cycle.

Delivery of light was controlled by a Master 8 controller. Seizure induction was attempted by the delivery of 10 Hz, 25 ms pulses of 473 nm light (Laserglow). Threshold duration and laser power were characterized in Figure S2A and S2B. Stimulation was performed every 1 – 3 hours in animals over the course of many days. After stimulation ceased, we performed post stimulation recording. Animals with no light induced response at all – not even a stimulation artifact – were excluded from the analysis.

### Pharmaceuticals

On the day of pharmaceutical testing, a few optical activations occurred prior to drug injection to determine the pre-drug induction success rate. Then, animals received either one subcutaneous injection of 5 mg/kg diazepam *(58)* (Dash Pharmaceuticals) or an intraperitoneal injection of 800 mg/kg levetiracetam *(55)* (Sigma Aldrich). Up to six more activations occurred every 1 – 1.5 hours following ASM administration. After the experiment, animals had a break of one to two days during which there was no optical stimulation. The type of ASM and order of ASM injection was assigned by recording cage position. Pharmacologic induction success rate and behavioral manifestation rate was compared using a one tailed Wilcoxon matched pairs signed rank test.

### Perfusion

After the recordings were completed, animals were deeply anesthetized and transcardially perfused with phosphate buffered saline (PBS) and then 4% paraformaldehyde in PBS. Brains were fixed overnight in 4% paraformaldehyde in PBS and then placed into 30% sucrose in PBS for cryoprotection. Once the brains have sunk, they were frozen in dry ice. Brains were sliced at 50-micron thickness in the coronal plane on a microtome and slices were stored in PBS with 0.05% sodium azide.

### Histology

Slices were either immediately visualized after slicing or underwent Nissl staining. Slices for immediate visualization were placed into a glass petri dish and directly imaged on a Leica DMIRB inverted fluorescence microscope using the LasX software. Other slices were stained with the Nissl staining protocol using 0.1% cresyl violet (Sigma Aldrich). Briefly, slices were dehydrated in increasing concentrations of reagent alcohol, beginning with 75%, then progressing to 95%, and ending with 100% reagent alcohol. Next, slices were demyelinated with xylene (Sigma Aldrich). After demyelination, slices were rehydrated with decreasing concentrations of reagent alcohol, beginning with 100%, then progressing to 95%, 75%, and 50% reagent alcohol. The final step of the rehydration is an incubation in DI water. After rehydrating, slices were stained in 0.1% cresyl violet in DI water for 20 to 40 minutes at room temperature. Slices were then rinsed in DI water, before being dehydrated and destained with fresh 50%, 75%, 95%, and 100% reagent alcohol. For the final cleaning, slices were rinsed in xylene. DPX mounting media (Sigma Aldrich) was used to mount the slices and the stained slices were imaged and stitched on a Leica microscope using the LasX software.

### Statistics and Computational Data Processing

#### A. Preprocessing

Recorded EEG data was filtered with second order IIR notch filter to remove 60 Hz line noise and then z-score normalized to a baseline period – defined as 5 seconds immediately preceding the onset of the optogenetic stimulus. A 4 Hz $6^{th}$ order Butterworth high pass filter was used to remove low frequency artifacts.

#### B. Feature Calculation

The following features were extracted from 500 millisecond windows with 250 millisecond displacement. In the below formulas, *N* represents the number of items in the recording, indexed by *n*, and *x* is an element in the recording.

1. Line length *(59)*, which measures the complexity of the signal: $\sum_{n=2}^{N} |x_n - x_{n-1}|$
2. Area *(60)* $\sum_{n=1}^{N} |x_n|$
3. Energy $\sum_{n=1}^{N} x_n^2$
4. Zero crossings around mean *(61)* $\sum_{n=2}^{N} \mathbf{1}\big((x_{n-1} - \bar{x} > 0 \wedge x_{n-1} - \bar{x} < 0)\big|(x_{n-1} - \bar{x} < 0 \wedge x_{n-1} - \bar{x} > 0)\big)$
5. Root mean squared amplitude $\sqrt{\frac{1}{N}\sum_{n=1}^{N} |x_n|^2}$
6. Skewness *(62)* $\frac{\frac{1}{N}\sum_{n=1}^{N}(x_n - \bar{x})^3}{\left(\sqrt{\frac{1}{N}\sum_{n=1}^{N}(x_n - \bar{x})^2}\right)^3}$
7. Approximate entropy using the MATLAB approximateEntropy function.
8. Lyapunov exponent, a measure of divergence, using the MATLAB lyapunovExponent function.
9. Phase locked high gamma *(63)*. Data was filtered into LFP (4 – 30 Hz) and high gamma (80 – 150 Hz) portions using a second order Butterworth filter. A Hilbert transform (H) was used to extract imaginary and real components from both LFP (*lfp*) and high gamma (*hg*) data. The LFP phase ϕ was calculated using the 2-argument archtangent. The high gamma amplitude (A) was calculated by taking the square root of the sum of the following: 1) the square of the imaginary component of the high gamma data and 2) the square of the real component of the high gamma data. The phase locked high gamma is calculated as the absolute value of the mean of the following product: $|\mathrm{H}(hg)| \times e^{i(\phi(lfp) - \mathrm{A}(hg))}$.
10. Magnitude squared coherence between channels *(64)* was calculated using the MATLAB mscohere function. Channels to calculate coherence with were defined to be between the set of hippocampal wires (Channel 1 and 2), the two EEG screws (Channel 3 and 4), the ipsilateral screw and wire (Channel 1 and 3), and the contralateral screw and wire (Channel 1 and 4). The function calculated the power spectral density (P) using the Welch's overlapped periodogram function. The function also calculated the cross power spectral density of the two channels' data (x). As an example, the magnitude squared coherence (C) for channel 1 and 2 was $\mathrm{C}_{1,2}(x) = \frac{|\mathrm{P}_{1,2}(x)|^2}{\mathrm{P}_{1,1}(x)\mathrm{P}_{2,2}(x)}$
11. Mean absolute deviation $\frac{1}{N}\sum_{n=1}^{N} |x_n - \bar{x}|$
12. Band power between 1 – 30 Hz, 30 – 300 Hz, and 300+ Hz. Band power was calculated using the MATLAB bandpower function, The function generated a power spectral density estimate using the Hamming window and integrated the area from the periodogram between the frequencies of interest.

After calculation, individual features were z – scored normalized by subtracting the mean of the feature values for the entire sequence from the feature values and dividing by the standard deviation of the feature values for the entire sequence.

#### C. Induced Activity Length Determination with K Nearest Neighbor Model

An unsupervised K Nearest Neighbor Model was trained by feeding in z-score normalized features from one spontaneous seizure (event #49 from animal 100) as the input. The training response variable was 3 distinct clusters identified by a K means model using the same input feature set. Two of the clusters were defined as epileptic, and the remaining cluster was defined as the baseline. Despite differences in each event's EEG morphology, about 80% of the time, the prediction for time of termination was within 5 seconds of the true event length. The other 20 % of the time, the termination time was manually adjusted to create a 'master' event duration list. The 'master' event duration was subsequently used for segregating the windows from the EEG feature space into the following categories: 1) before stimulation (5 seconds), 2) during stimulation, 3) initial/beginning third of the induced activity, 4) second/middle third of the induced activity, 5) final/ending third of the induced activity, and 6) post stimulation (30 seconds). Event durations were averaged per animal, and a two tailed Wilcoxon rank sum test was used to compare average of the naïve animals to the average of the epileptic animals. Overall induction success rate was also compared with a two tailed Wilcoxon rank sum test between naïve and epileptic animals. However, daily activation success rate (by day since first stimulation) was compared using a two sided pairwise t test.

#### D. Threshold Determination

The threshold for activity induction was defined as the minimum duration and laser power at which 10 Hz optical stimulus would cause a minimum five-second-long electrographic activity at least 66.67% of the time. Only drug-free conditions were considered. A two tailed Wilcoxon rank sum test was used to compare the threshold duration and the threshold power distribution of the naïve and epileptic animals.

#### E. Spontaneous Seizure Detection in Freely Moving Animals

Spontaneous seizures were detected using custom-written Matlab code *(56)*. Briefly, three EEG features involving median, line length, and the difference between local maximum and minimum were calculated for each time bin for all four electrodes, and time points at which the feature value exceeded a threshold were listed as seizure candidate time points. Using custom EEG reviewing software, each seizure candidate time point was visually inspected by an expert reviewer, and behavioral seizures were verified through the video file. Daily spontaneous seizure frequency in epileptic animals before and after stimulation began was compared using a two tailed Mann Whitney test. Daily spontaneous seizure frequency was compared between CA1 IHK and CA3 IHK group using a two tailed Mann Whitney test.

#### F. Video EEG Behavioral Scoring

Video EEG of each event was manually reviewed twice by an expert reviewer. Uncontrolled behaviors and the Racine scale *(57)* associated with the behaviors are as follows in Table S1:

**Table S1 Behavioral Scoring and Racine Scale Assignments**

| **Racine** | **Behaviors** |
|---|---|
| **0** | Normal behavior, grooming, free movement (even with slight limp) |
| **1** | Clear Freezing / Flattening of the Body |
| **2** | Tail Stiffening / Forelimb Clonus / Uncontrolled Shaking |
| **3** | Rearing |
| **4** | Wild Run / Backpedaling |
| **5** | Uncontrolled Jumping / Loss of Righting Reflex |

The Racine behavior score of the event was determined to be the highest value out of all the behaviors the animal displayed during each event. If there was no uncontrolled behavior displayed, or if the animal did not move at all throughout the entire event, the score was 0 – no change.

The mean Racine level of induced activity was compared between naïve and epileptic animals by a two sided pairwise t test. A two sided pairwise t test was also used to compare the rate of electrographic events that had a behavioral manifestation (a score above 0).

#### G. Linear Mixed Effect Models

Mixed effects models were fitted with the lmer function in the lmerTest package in R. Feature values calculated in Section B and segregated in Section C were exported from Matlab to R. The baseline was set to be the pre-stimulation values for each feature. Since we wanted to compare the extent to which features changed from pre-stimulation values to each of the event terciles, the time point was a fixed effect in all mixed effect models. The animal was set as the random effect due to differences in the induced activity between animals.

The other fixed effect in the mixed effect model depended on what groups we were comparing. Between the epileptic and naïve animals, the fixed effect was whether the animal was epileptic or not. Between the spontaneous seizures and induced activity, the fixed effect was whether there was an optogenetic stimulus. The mixed effect model equation was thus: EEG.Feature ~ Comparison * Time.Point + (1 | Animal).

As mentioned in the main text, additional filtering of events occurred before they were fed into the mixed effect model. Naïve events differed significantly between early and late activation days, so for the epileptic versus naïve induction comparisons, events were split by the day since stimulation start (day 1 – 4 and day 5 +). The spontaneous seizure analyzer (Section E) could only identify behavioral seizures with Racine score 3 or above. Thus, for the spontaneous seizures versus induced activity comparison in epileptic animals, only events with Racine score of 3 or above were compared. For both comparisons, events shorter than 15 seconds were excluded to reduce noise from shorter afterdischarges. All drug treatment trials were also removed.

#### H. Support Vector Machine Classification of Spontaneous and Induced Activity

A new support vector machine for classifying induced activity was trained per animal. First, features were calculated from ten baseline EEG segments from before stimulation began. Next, these features were combined with the features from the animal's spontaneous seizures to form the training inputs. The SVM algorithm was trained using that input set on separating events as either 'spontaneous seizure' or 'baseline'. The testing data consisted of features from all optical activations. The model was asked to classify the induced activity. Accuracy and error rates were calculated based on ground truth, where a successful induction was defined as an electrographic event with afterdischarges lasting a minimum of 5 seconds.

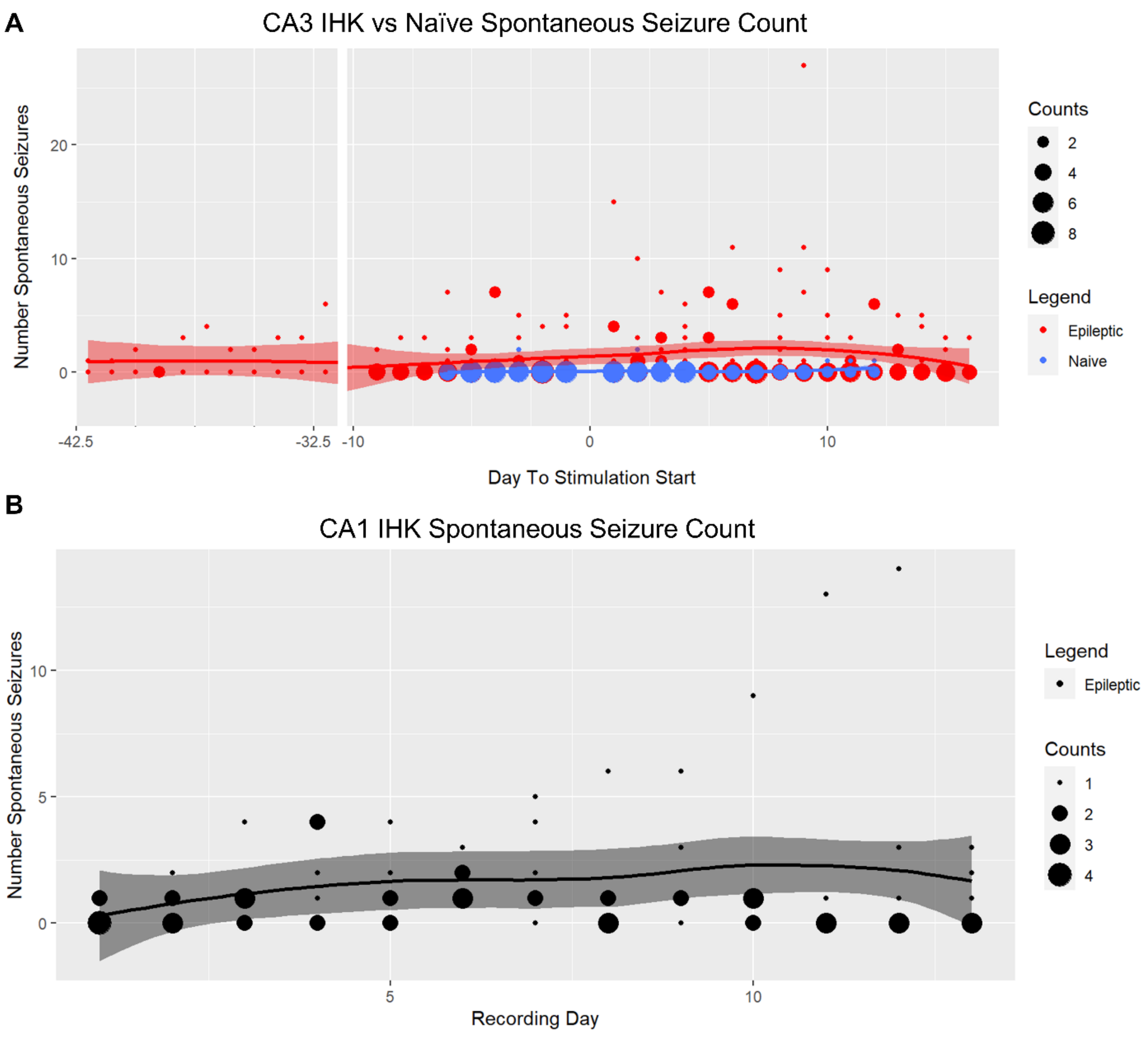


**Figure S1: Spontaneous seizure counts in freely moving animals.** (**A**) Average daily spontaneous seizure counts in ten CA3 IHK epileptic animals was 1.7 spontaneous seizures per day. Average daily spontaneous seizure frequency slightly increased after stimulation began (1.8 after vs 0.98 before) but the difference was not statistically significant ($p = 0.6354$, Mann Whitney Test). After optical stimulation, 3 naïve animals began experiencing spontaneous seizures. One naïve animal (out of 7) experienced spontaneous seizures prior to stimulation. (**B**) Average daily spontaneous seizures counts in 6 CA1 IHK epileptic animals was 1.6 spontaneous seizures per day.

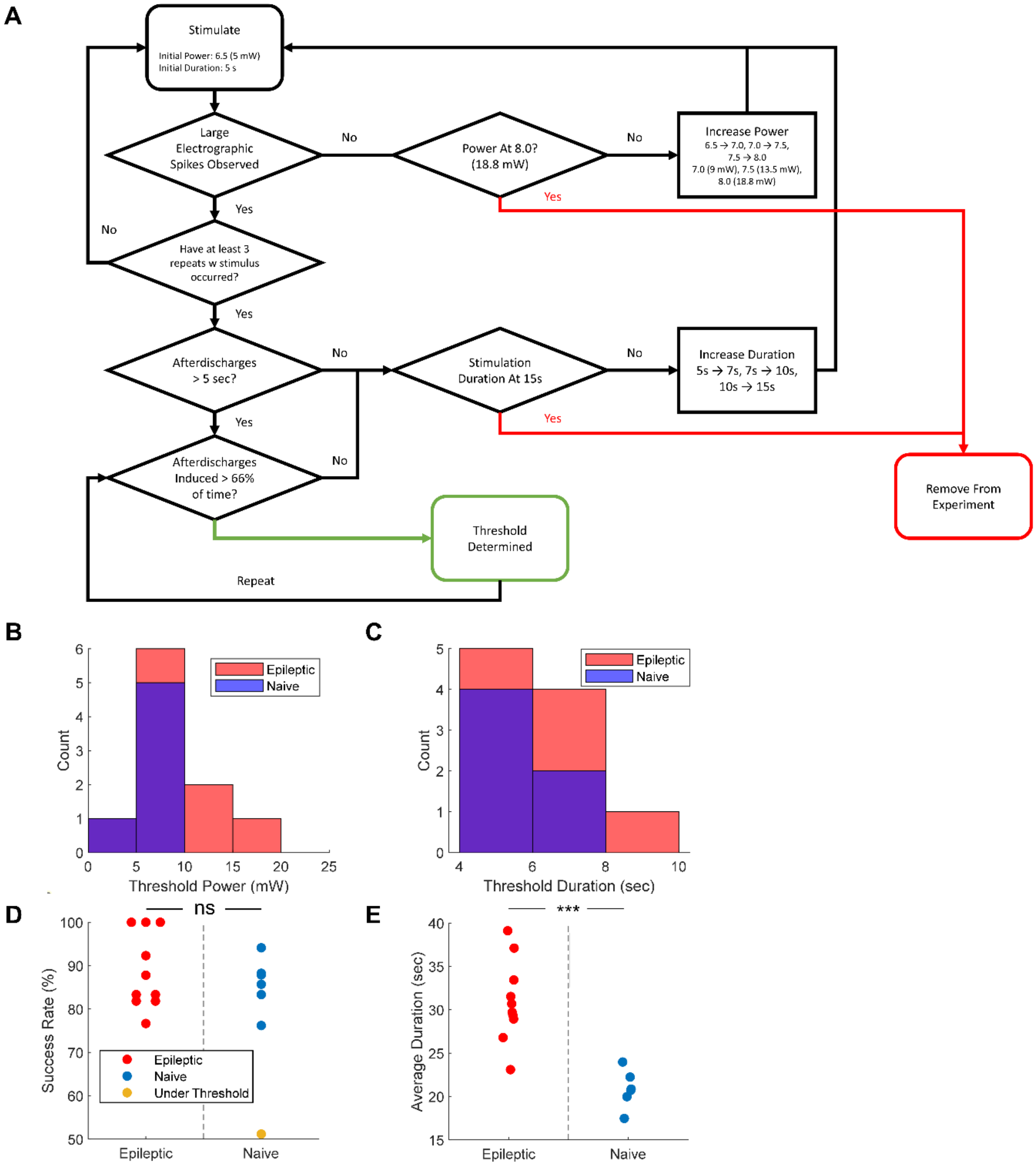


**Figure S2: Threshold for effective stimulation in freely moving animals.** **(A)** Flowchart for determination of threshold stimulus. Threshold stimulus's effect is continuously monitored throughout the experiment and adjusted as necessary. **(B)** Threshold power to induce electrographic activity for n = 10 epileptic (9.14 ± 4.75 mW) and n = 6 naïve animals (6.17 ± 1.58 mW) (Wilcoxon rank sum test, $p = 0.137$). **(C)** Threshold duration was comparable between the same epileptic animals (6.30 ± 1.64 s) and naïve animals (5.67 ± 1.03 s) (Wilcoxon rank sum test, $p = 0.7133$). **(D)** Epileptic animals had comparable success rate of induction (88% ± 8.8 %) at threshold to naïve animals (86 ± 6.0 %) (Wilcoxon rank sum test, $p = 0.8506$). One naïve animal never reached consistent induction above 66.7%. **(E)** With the threshold stimulus, induced activity mean duration was longer in epileptic animals (30.98 ± 4.69 s) than in naïve animals (20.87 ± 2.19 s) (Wilcoxon rank sum test, $p = 0.0005$).

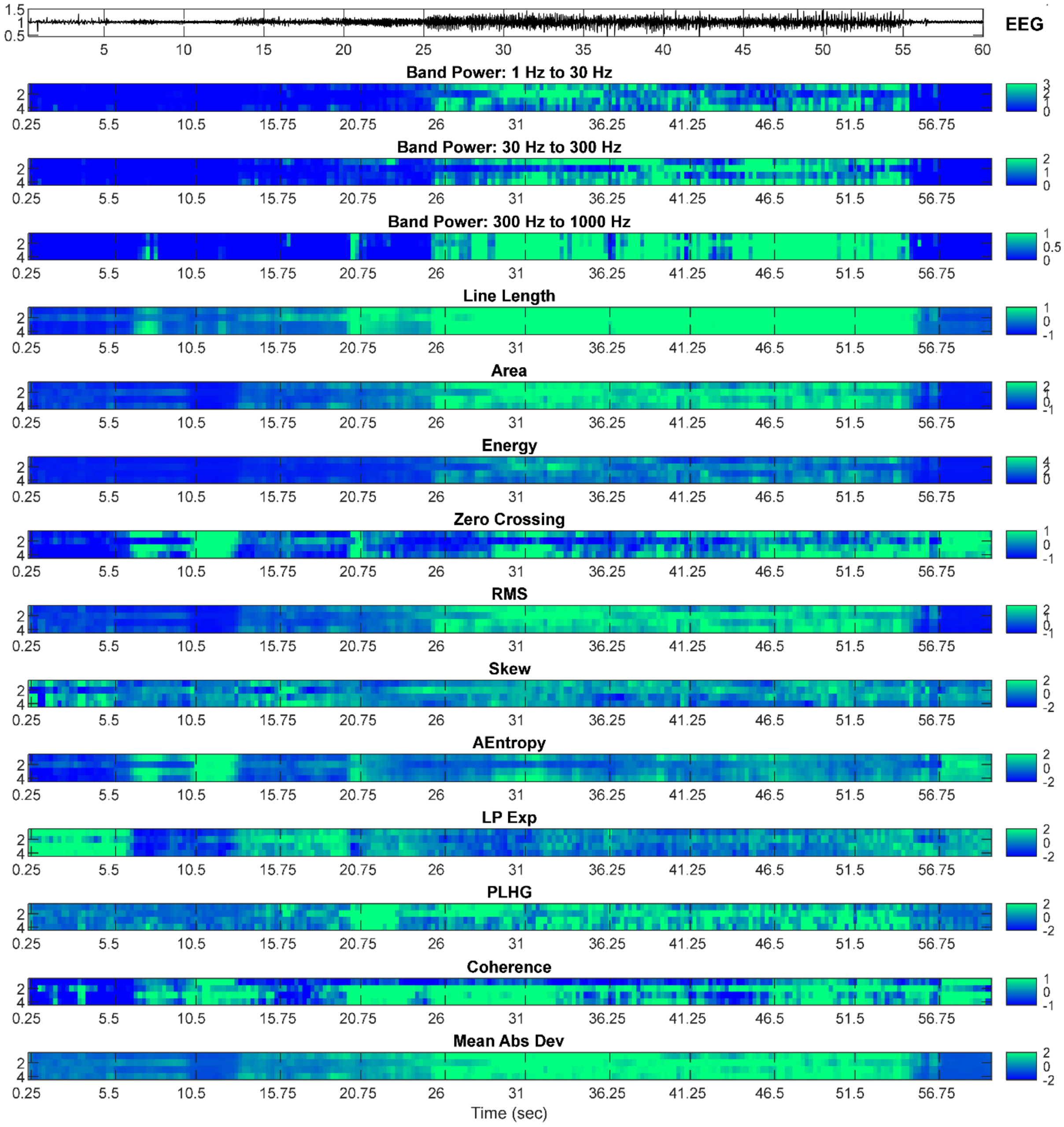


**Figure S3: For each channel, numerous EEG features were calculated and z-score normalized for feeding into K Nearest Neighbor Classifier** (Fig. S4). Feature output calculated for the induced seizure in Fig. 3B. RMS – Root Mean Squared Amplitude. AEntropy – Approximate Entropy. LP Exp – Lyapunov Exponent. PLHG – Phase Locked High Gamma. Mean Abs Dev – Mean Absolute Deviation.

.

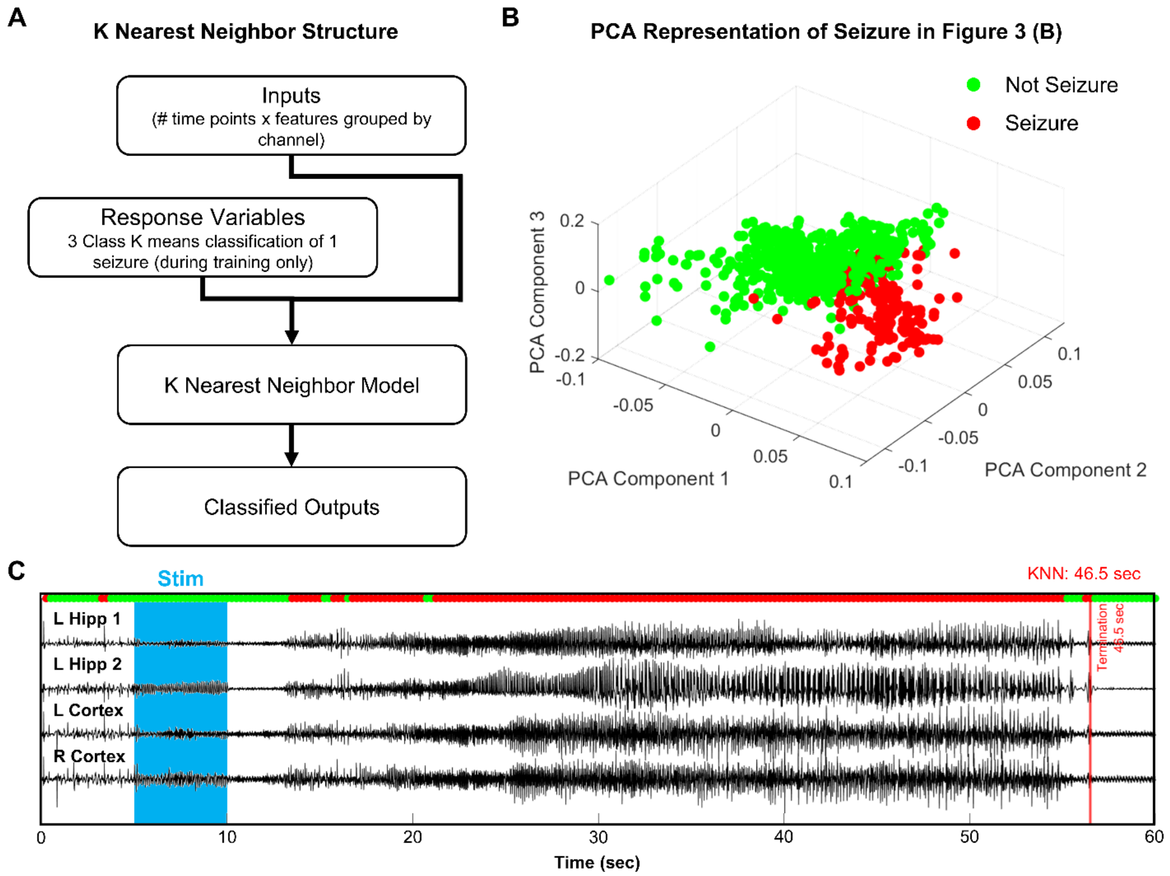


**Figure S4: Three class k-nearest neighbor (KNN) model predicted activity duration.** **(A)** Model was trained using Z scored normalized EEG features from one spontaneous seizure. **(B)** 3-dimensional PCA space distribution of computationally determined seizure/not seizure classes for seizure depicted in Fig. 3B. **(C)** KNN predictions plotted against filtered EEG waveforms for seizure depicted in Fig. 3B. Model determined that activity lasted for 46.5 s after stimulation terminated.

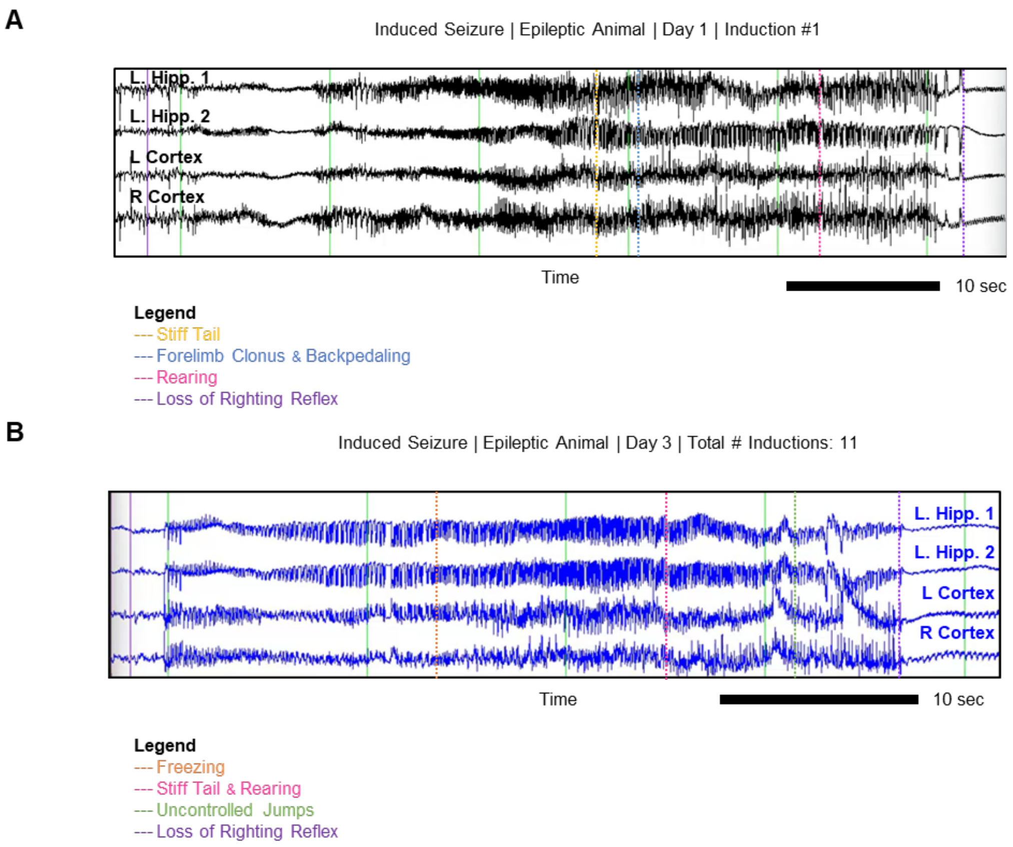


**Figure S5: Examples of Racine 5 behavioral and electrographic seizures induced in epileptic animals.** **(A)** Seizure depicted in Figure 3(B), from animal #100. Key frames depict stiffening of the tail (yellow), forelimb clonus with backpedaling (blue), rearing (pink), and loss of righting reflex (purple), with approximate locations of the frame marked by lines on the waveform. **(B)** Example of induced seizure from another epileptic animal (#107). In this seizure, onset was characterized by freezing (orange) before progressing to clonus, tail stiffening, rearing (pink) and – eventually – uncontrolled jumps (green) and loss of righting reflex (purple).

.

Induced Seizure | Epileptic Animal | Day 1 | Induction #3

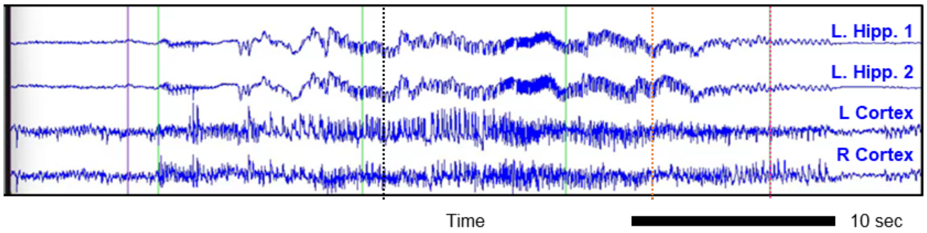


**Legend**
--- Independent Movement
--- Freezing
--- Stiff Tail

**Figure S6: Example of Racine 2 behavioral and electrographic seizure induced in epileptic animal #107.** Animal shown here is the same as the one in Fig. S5B. Stimulus used in this evocation was lower in intensity and duration than the threshold stimulus for consistent seizure evocation. Approximate locations of key frames marked in color. Initially, there was no obvious seizure behavior (black). However, the animal froze (orange) and the tail stiffened (pink) towards the end of the event, signaling a seizure has occurred.

Failed Induction | Epileptic Animal | Day 1 | Induction #2

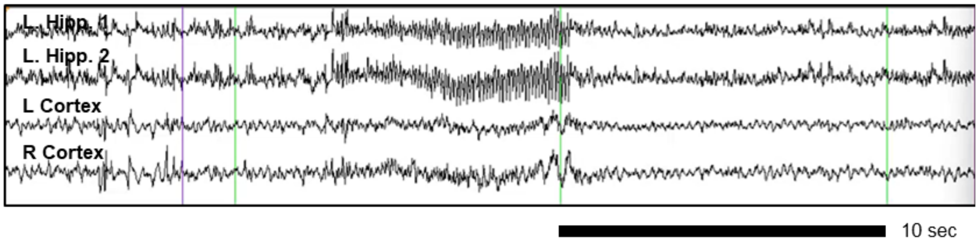


**Figure S7: Example of failed induction in epileptic animal (#105).** Stimulus elicited no electrographic discharges. In this trial, stimulus power was under that of the threshold stimulus for consistent induction in this animal.

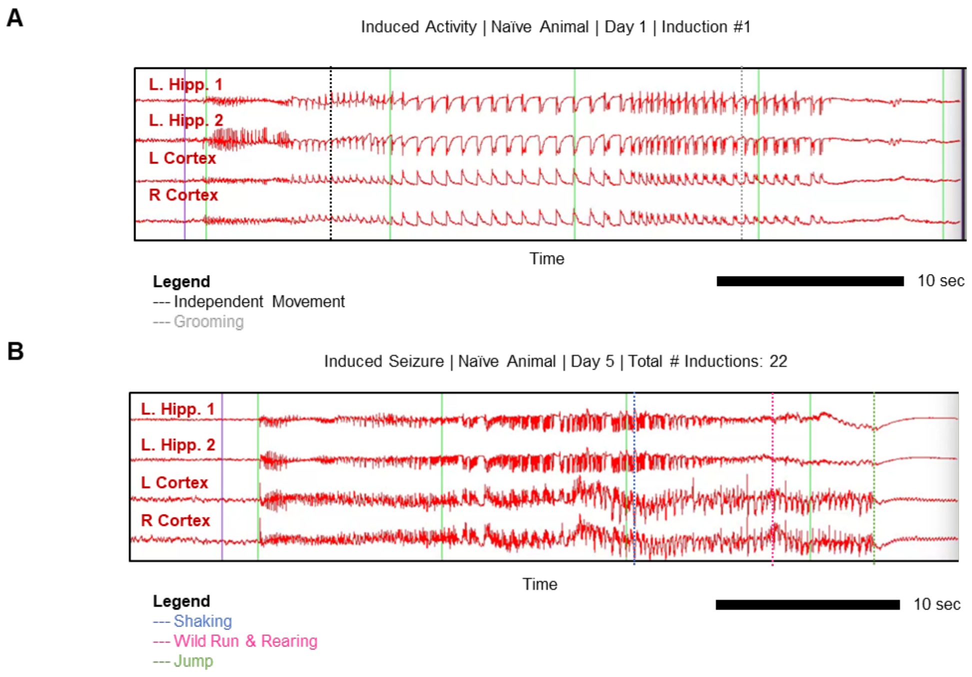


**Figure S8: Examples of inductions in naïve animal #110. (A)** Afterdischarges depicted in Figure 4(A). Animal displayed independent movement (black) and grooming (gray) during spiking. Behavioral Score: No seizure. Electrographic Score: Afterdischarges induced. **(B)** Seizure depicted in Figure 4(B) Animal begins shaking (blue) before embarking on a wild run with a stiff tail (pink). Finally, the animal ended up rearing and performing an uncontrolled jump (purple). Behavioral Score: Racine 5. Electrographic Score: Seizure induced.

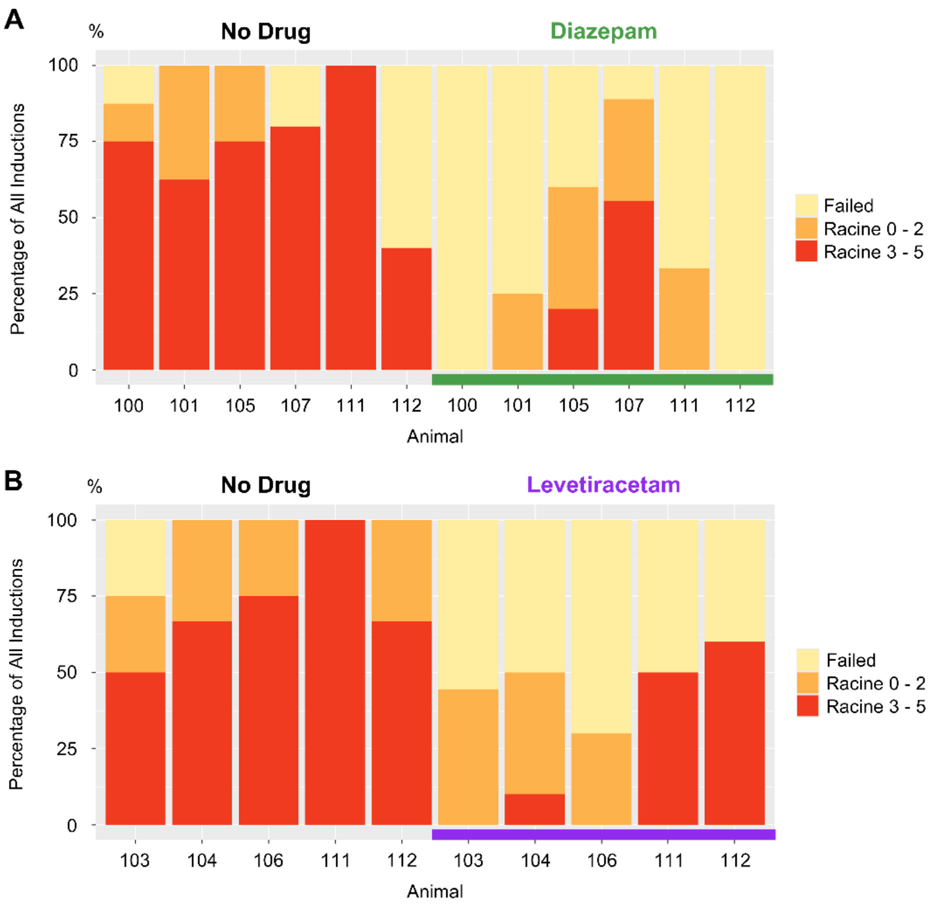


**Figure S9: Racine level of seizures, measured as percentage of all inductions in epileptic animals before & after drug application. (A)** 5 mg/kg Diazepam reduces the Racine level of behavioral seizures in all animals, including animal 107, which was a diazepam non-responder on one of two testing days. **(B)** Levetiracetam also reduces the Racine level of behavioral seizures in all animals. In both graphs, data between multiple testing days was aggregated by animal.

A

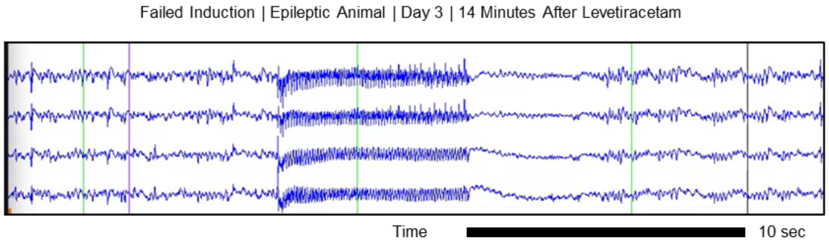


B

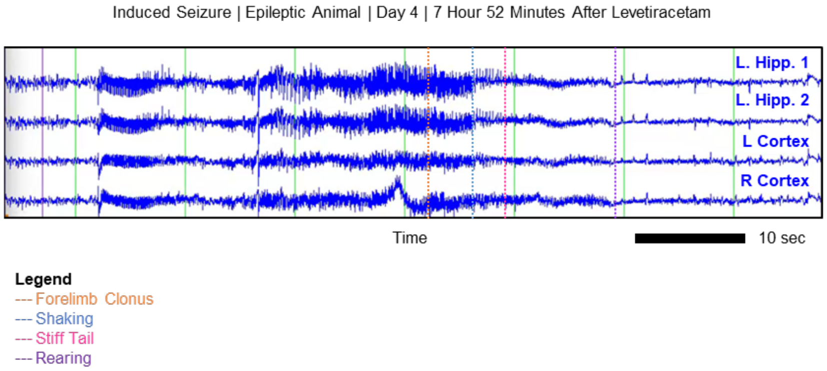


**Figure S10: Inductions after intraperitoneal (IP) injection of 800 mg/kg levetiracetam. (A)** In animal #104, electrographic activity could not be induced by the threshold stimulus 14 minutes after levetiracetam administration. Behavioral Score: No seizure. Electrographic Score: Failed. **(B)** In the same animal, 7 hours and 52 minutes after levetiracetam administration, behavioral seizures, characterized by clonus (orange), shaking (blue), tail stiffening (pink) and rearing (purple) could be induced. Behavioral Score: 3. Electrographic Score: Seizure Induced.
